# GenomeProt: User friendly proteogenomics for canonical and non-canonical proteoform characterisation

**DOI:** 10.64898/2026.08.06.743133

**Authors:** Hitesh Kore, Josie Gleeson, Ching Yin Wan, Ricardo De Paoli-Iseppi, Mriga Dutt, Yair D.J. Prawer, Arwa Alkaraki, Andrew Lonsdale, Christine A. Wells, Lorey Smith, Michael B. Clark, Benjamin L. Parker

## Abstract

Quantifying the diversity of RNAs and proteins produced by cells is fundamental to the biological and clinical sciences. However, many RNAs and proteins remain uncharacterised, especially proteins translated from alternate RNA isoforms; untranslated regions of mRNAs and non-coding RNAs, as well as the effects of DNA variation on protein sequences. Proteogenomics aims to characterise the complete proteome by integrating genomics and/or transcriptomics with proteomics, but current tools have limitations in useability, analysis features and visualisation of resulting data. To address these gaps, we developed GenomeProt, a user-friendly GUI-based tool for integrative proteogenomic analysis. We demonstrate its utility by integrating long-read RNA sequencing with mass-spectrometry-based proteomics to pinpoint proteoform expression generated by alternative splicing; discover novel, unannotated proteins in human brain samples; and quantify variant-containing peptides associated with treatment resistance in a melanoma xenograft model. GenomeProt brings the discovery power of proteogenomics to biologists, illuminating the hidden proteome.

## INTRODUCTION

The accurate and comprehensive characterisation of the products encoded by the genome is a fundamental requirement to understanding gene functions and organism-level health and disease. Investigation of the RNAs transcribed by the genome with techniques such as RNA-seq has revealed vast numbers of alternative products (RNA isoforms) made by genes, as well as non-coding genes that appear not to encode proteins (1, 2). Simultaneously, application of techniques such as Ribo-seq and mass-spectrometry have dramatically expanded our understanding of the translated products made from these RNAs (3). These have identified wide-spread translation of: i) alternative RNA isoforms into alternative proteoforms; ii) non-canonical open reading frames (ncORFs), such as proteins/peptides from 5′ and 3′ untranslated regions (UTRs) of messenger RNAs (mRNAs) and currently annotated non-coding RNAs (ncRNAs); and iii) alternative protein variants due to inherited or somatic genetic variations that alter the protein sequence. However, despite these advances, our understanding of the RNA and protein products made by cells is far from complete, even in heavily studies species such as humans.

A key limitation to characterising novel translated products has been the standard application of mass spectrometry-based proteomics, which compares the peptide spectrum from a sample to standard reference databases such as UniProtKB (Swiss-Prot/TrEMBL) or NCBI (RefSeq/Entrez). In this method, only proteins present in the database can be identified. While these databases aim to contain all common protein-coding sequences in the genome, they may not contain alternative proteoforms; ncORFs; proteins from novel genes; or proteins containing amino acid variants. Furthermore, these databases contain proteins/proteoforms that are not expressed in the sample(s) being analysed, leading to ambiguous protein and proteoform assignments when identified peptides also match non-expressed database entries.

Proteogenomics has been developed to overcome the limitations of standard mass-spectrometry-based proteomics for protein identification. Proteogenomics uses genomics and/or transcriptomics to generate customised *in silico* databases of predicted ORFs that are used to analyse mass spectrometry-based proteomic data (4). Use of sample-specific databases therefore enables investigators to discover novel proteins/translated products and more accurately pin-point proteoform expression. Recent advances in proteogenomic methodologies have incorporated long-read RNA-sequencing based transcriptomics approaches to better define the RNA isoforms present in the sample and improve the accuracy of the ORF database (5, 6)

One of the most comprehensive proteogenomic studies for the characterisation of alternative splicing and proteoform expression was recently presented by Sinitcyn et al., who provided direct evidence that hundreds of predicted alternatively spliced isoforms are translated (7). Proteogenomic studies have also revealed extensive translation of ncORFs outside of annotated protein-coding regions of mRNAs (8), including 7,264 previously unannotated ncORFs identified by the TransCODE consortium following proteogenomic analysis of 95,520 proteomics experiments (9) while functional analyses have confirmed roles for ncORFs in human physiology and disease (10, 11). In addition, emerging evidence suggests microproteins from ncORFs can also be presented on Human Leukocyte Antigen (HLA) molecules indicating their potential relevance for immunotherapeutic strategies (12–15).

Proteogenomics can also be used to investigate the impact of genetic variants on protein sequences. The Clinical Proteomic Tumour Analysis Consortium (CPTAC) pioneered approaches for characterising the effects of patient-specific genetic variants across tumours including colon, rectal, breast, ovarian and renal cancers (16–19). More recently, the impact of patient germline variants and rare pathogenic variants on proteoform function has also been studied across multiple cancer types (20). Taken together, these studies highlight the impact of proteogenomics in discovering; i) alternatively spliced transcripts that produce multiple proteoforms each with distinct function, ii) novel proteins from ncORFs that may play important roles in biology and/or represent new therapeutic/diagnostic targets, and iii) specific variants or mutations that impact protein sequence and potentially protein function.

Despite the power of proteogenomics for genome characterisation and its wide relevance to researchers across the biological and clinical sciences, performing proteogenomic analyses remains challenging. While there are several computational packages for proteogenomics (for a reviews see (21, 22)), a common theme is they require dedicated bioinformatics experience; are not widely applicable to different types of RNA-seq data; don’t enable user-friendly interrogation and visualisation of peptide mappings to the genome/transcriptome for proteoform characterisation; and the databases and integrated outputs generated may lack comprehensive proteome or genome annotations, which are important for downstream analyses. To address these limitations and improve the accessibility of proteogenomics we developed GenomeProt, a user-friendly, graphical user interface (GUI)-based tool that can perform proteogenomic analyses from both short-read and long-read RNA-seq data, with or without the inclusion of genetic variants. We applied GenomeProt to several projects combining paired Oxford Nanopore long-read RNA-seq with deep mass-spectrometry datasets generated using multiple proteases. We validated proteoform generation from alternative splicing in an *in vitro* cell line and identified thousands of candidate novel proteins in human brain. Finally, we showcase the use of GenomeProt to identify coding variants arising during melanoma treatment resistance in a patient xenograft model that were identified from whole genome sequencing (WGS).

## RESULTS

### Overview of GenomeProt

GenomeProt is a user-friendly, GUI-based workflow designed to facilitate integrative proteogenomic analyses in four steps: i) RNA-seq derived proteome database generation, ii) proteomics analysis, iii) proteogenomics data integration, and iv) data analysis and visualisation **(Fig. 1a** and **Supplementary video)**. In the database generation step, users can input FASTQ, BAM, or GTF files derived from long-read sequencing platforms, including Oxford Nanopore and PacBio. GenomeProt also supports database generation from short-read RNA-seq data with transcript abundance estimation using salmon (23). However, for precise identification and quantification of isoforms, long-read RNA-seq is recommended. FASTQ files are aligned to a reference genome using minimap2 (24), and expressed isoforms identified using bambu (25). The resulting transcript sequences are translated into three reading frames, collapsed into non-redundant ORFs and filtered, to generate a custom proteome database (see methods). A metadata file containing comprehensive annotations for each predicted ORF is also generated. This includes mapping ORFs against RefSeq, UniProt and OpenProt (26) to identify canonical proteins. This information is used to generate detailed FASTA headers to enable user friendly downstream analysis and visualisation. In addition to canonical ORFs, GenomeProt annotates ncORFs such as upstream ORFs (uORFs), upstream overlapping ORFs (uoORFs), downstream ORFs (dORFs), internal ORFs (intORFs) and intergenic ORFs (igORFs) which includes non-coding ORFs (ncORFs) with no overlap to annotated protein-coding genes (27, 28).

**Figure 1:**
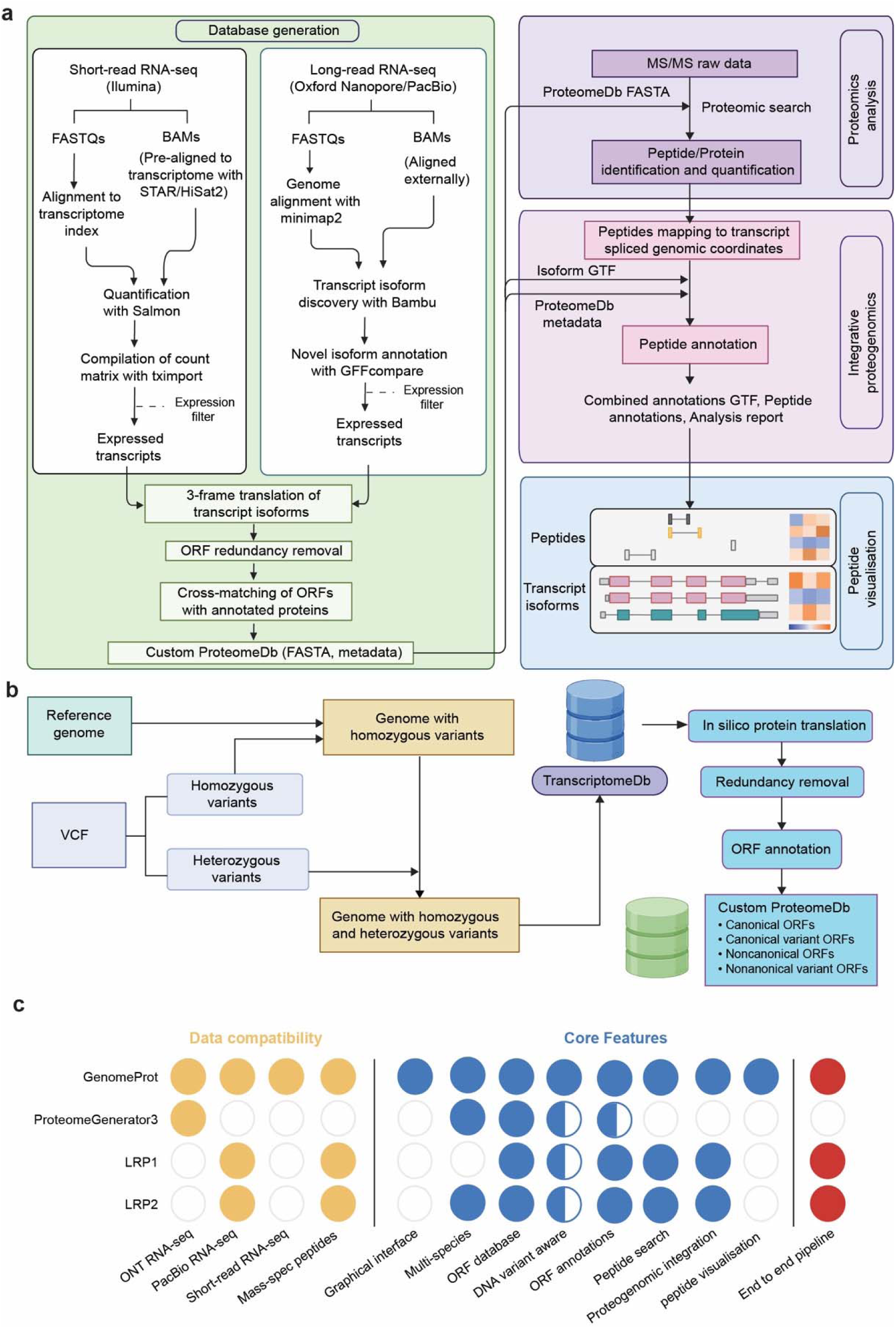
Schematic overview of GenomeProt workflow. (**a**) The workflow consists of four steps- (1) Database generation: A custom protein sequence database is constructed from transcriptomic data obtained using Short-read or Long-read RNA-seq. (2) Proteomic analysis: Tandem mass spectrometry (MS/MS) raw data are searched against the custom database for peptide and protein identification and quantification. (3) Integrative proteogenomics: Identified peptides are mapped to transcript coordinates, enabling annotation and discovery of canonical and non-canonical ORFs (ncORFs). (4) Visualisation: Peptides identified in step 3 are visualised on alternatively spliced transcripts or ncORF coordinates. (**b**) Overview of the variant aware module of GenomeProt incorporating mutations from variant call format (VCF) into RNAseq-derived proteome database. (**c**) Feature comparison of GenomeProt against proteogenomics pipelines.

GenomeProt also supports single nucleotide variant-aware proteome database generation using a multi-sample VCF file. The workflow produces two genome builds: one containing only homozygous variants and another containing both homozygous and heterozygous variants. Transcriptomes are reconstructed from each genome, followed by ORF prediction and redundancy removal, to produce a final variant-aware proteome database (**Fig. 1b**).

The proteomics analysis searches mass spectrometry data against the custom proteome database resulting in peptide and protein identifications. We incorporated FragPipe (29) within GenomeProt to enable this step. Users with an alternate preferred search engine such as MaxQuant, ProteomeDiscoverer, DIA-NN or Spectronaut, etc, can perform this step externally as GenomeProt provides standardised FASTA output for proteomics search engines and is compatible with their outputs.

The third step is proteogenomics data integration where peptides identified in the previous step are mapped to proteoforms and the resulting peptide and transcript coordinates are stored in a GTF file. The final step enables visualisation and analysis of peptide and transcript mappings onto the genome along with transcript and proteoform quantitative data. It allows gene-based queries and interactive visualisation of translated ORFs, the known and unannotated RNA isoforms that are translated, alongside the peptide evidence for translation.

GenomeProt is available in several forms to provide flexibility to meet user needs, these include a public webserver, user-installed GUI version and user-installed command line version.

### Comparison of GenomeProt to existing pipelines for proteogenomics

We assessed the features and usability of GenomeProt compared to other recent proteogenomic pipelines that are compatible with long-read RNA-seq. Specifically, work by Miller et al. (5), Kulej, et al. who presented ProteomeGenerator3 (PG3) (6) and Schertzer, et al. who recently developed LRP2 (30). **Supplementary Table S1 and Fig. 1c** compare how GenomeProt builds on these tools. Key features of GenomeProt include: i) enhanced usability due to its graphical user interface; ii) flexibility in input data types including both short-read and long-read RNA-seq; iii) applicability to multiple species including *Homo sapiens*, *Mus musculus*, *Rattus norvegicus*, *Drosophila melanogaster*, *Caenorhabditis elegans*, and *Danio rerio*, iv) extended metadata annotations, including peptide annotation to the transcriptome, identification of exon-junction-spanning peptides and the annotation of ncORFs, v) the ability to build proteome databases containing amino acid variants, and vi) an integrated visualisation module that displays the transcript isoforms, ORFs, peptides and proteins identified, and enables comparison of transcript and peptide intensities across experimental conditions. While previous tools have been developed to analyse/visualise proteomic data in the context of the genome and transcriptome (e.g. PG Nexus (31), PGx (32) and ProBAM (33)), performing the analysis requires use of the command line followed by visualisation with additional external software. GenomeProt was designed with the goal of improving the accessibility of proteogenomics and importantly, increasing the ability of researchers to perform downstream analysis and detailed characterisation of proteoforms.

### Assessing GenomeProt for proteoform discovery and splicing characterisation

To evaluate optimal run parameters for GenomeProt and maximise performance, we generated paired long-read RNA-seq data and tryptic/LysC peptide 2D-LC-MS/MS data of the Kusa 4B10 osteoblast cell line. The long-read RNA-seq data identified 36,400 non-redundant isoforms with a total read count >5, which were used to generate various proteogenomic databases. Our first goal was to compare peptide and protein identifications using canonical databases to GenomeProt databases using different ORF length parameters. Here we compared: i) Mouse UniProt canonical (∼21K entries) or UniProt canonical with isoforms (∼63K entries), ii) Mouse OpenProt (∼675K entries), and iii) GenomeProt generated databases from the long-read RNA-seq with various minimum ORF length cut-offs (in amino acids; >10aa, >30aa, >50aa and >100aa) with or without the inclusion of uORF and dORF >10aa (database sizes ∼32K – 291K entries) **(Supplementary Fig. 1a)**. Overall, the number of peptides and proteins identified from the various databases at 1% protein level FDR were highly similar despite differences in database sizes **(Supplementary Fig. 1b-c)**. The inclusion of annotated isoforms in UniProt or the use of OpenProt slightly increased the number of proteins identified compared to the UniProt canonical database (**Supplementary Fig. 1b-c**). However, the larger databases required slightly higher discriminate score (CScore) cut-offs to achieve 1% FDR (**Supplementary Fig. 1d**). Compared to the larger UniProt- and GenomeProt-generated databases, ∼3,600 peptides were identified from only the smaller UniProt canonical database and these were from 91 proteins that failed to pass the stricter CScore cut-off required for larger database searches (**Supplementary Fig. 1e)**. Conversely the GenomeProt databases also identified similar numbers of peptides not found in UniProt (see below for more detailed analysis), confirming the potential of the pipeline to identify peptides not found in reference databases. This highlights the potential of GenomeProt for proteoform discovery.

We next focused our analysis on the results obtained using the GenomeProt database containing ORFs >30aa length with the inclusion of uORFs >10aa in length (∼134K entries). The GenomeProt database resulted in a greater number of peptides that map to only a single entry in the database, as well as more unique proteotypic peptides (∼70K in GenomeProt verses, ∼54K in UniProt with isoforms verses ∼11K in OpenProt), although the total number of peptides and proteins identified were similar to UniProt/OpenProt (**Supplementary Fig. 1f**). To provide a more comprehensive characterisation for downstream analysis, we performed additional 2D-LC-MS/MS via digestion with AspN or GluC. The trypsin/LysC, AspN and GluC data were searched against both the GenomeProt database containing ORFs with >30aa length (including uORFs >10 aa; ∼134K entries), and the UniProt canonical with isoforms databases (∼63K entries). The combined results from the multiple protease analysis revealed that the GenomeProt database identified 245,904 peptide sequences on 12,580 proteins groups while the UniProt database identified 257,942 peptide sequences on 12,399 protein groups (**Supplementary Table S2**). Importantly, GenomeProt identified a greater number of uniquely mapped proteotypic peptides (101,484 in GenomeProt [41%] verses 83,959 in UniProt [32%]) (**Fig. 2a**). As a result, there were 37,846 peptides mapping to a single proteoform in the GenomeProt-derived database that were mapped to multiple proteoforms in UniProt (**Fig 2b**). These results suggest that GenomeProt can more accurately discriminate the expressed proteoforms.

**Figure 2:**
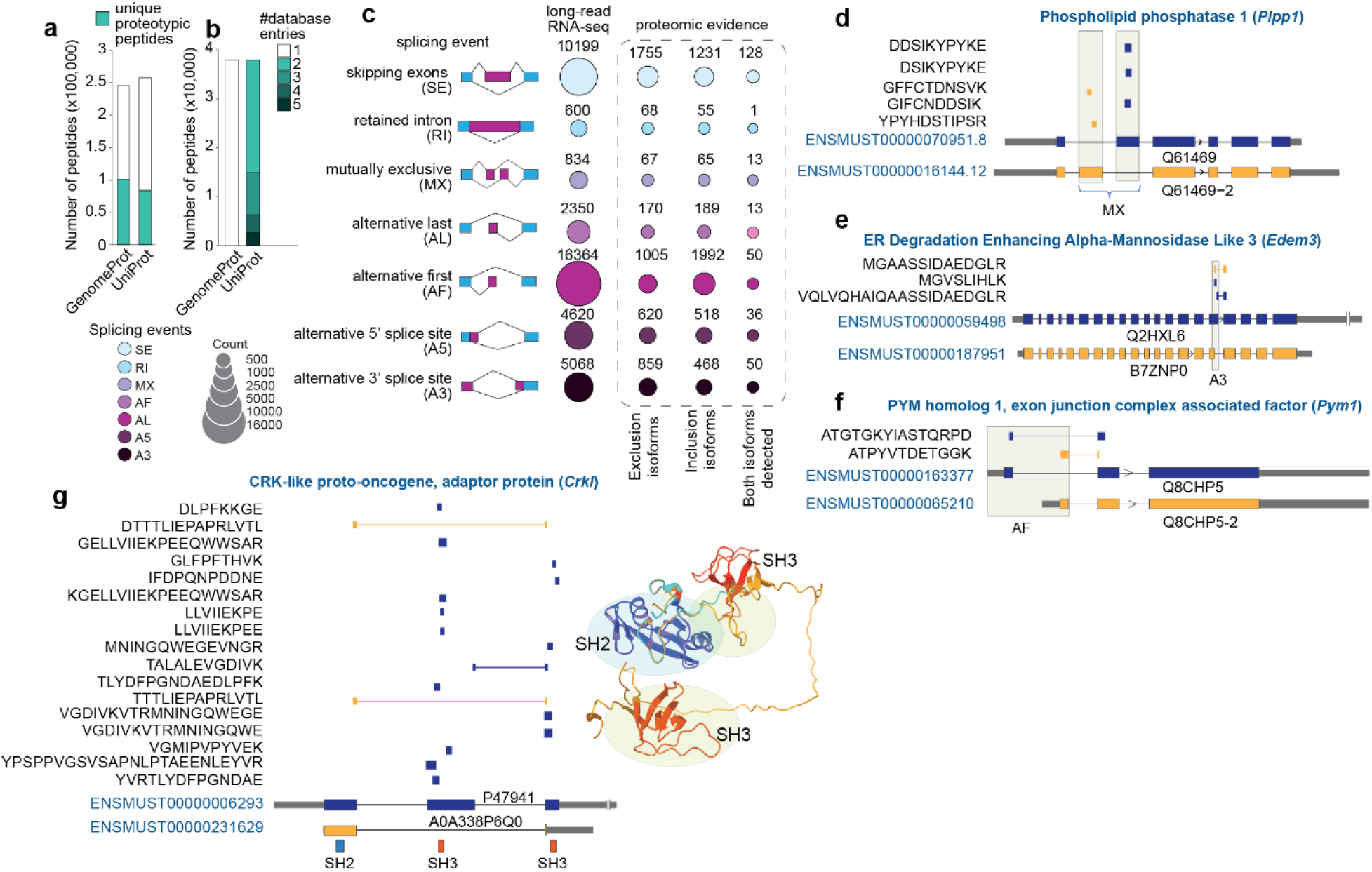
Evaluation of the GenomeProt database and identification of alternative splicing events in mouse osteoblast Kusa cells. (**a**) Number of proteotypic and shared peptides identified using the GenomeProt and UniProt databases. (**b**) Number of GenomeProt proteotypic peptides that mapped to multiple proteoforms in the UniProt database. (**c**) Alternative splicing events identified by long-read RNA sequencing and those supported by proteomic evidence. Bubble size indicates the number of events detected for each splicing category, including skipped exon (SE), retained intron (RI), mutually exclusive exon (MX), alternative last exon (AL), alternative first exon (AF), alternative 5′ splice site (A5) and alternative 3′ splice site (A3). Numbers indicate the total number of events identified by each modality. Exclusion transcripts: exon skipped isoforms in SE; fully spliced isoforms in RI; short isoforms in A3 and A5; isoforms with 3′ MX exon. Inclusion transcripts: exon included isoforms in SE; intron retained isoforms in RI; long isoforms in A3 and A5; isoforms with 5′ MX exon. Representative examples of alternative splicing events in *Plpp1* (**d**), *Edem3* (**e**) and *Pym1* (**f**), corresponding to mutually exclusive exon (MX), alternative 3′ splice site (A3) and alternative first exon (AF) events, respectively. Alternative exons and splice junctions are supported by multiple proteotypic peptides. Blue and orange represent alternative isoforms, and identified peptides are shown above each gene model. (**g**) Alternative splicing of *Crkl* produced isoforms with distinct domain architectures. Proteotypic peptides provide proteomic evidence for their expression. Alternative isoform A0A338P6Q alters the canonical domain organisation, as shown by predicted protein structures highlighting changes in SH2 (14–88 aa; blue) and SH3 domain arrangement (130–175 aa, 239–293 aa; orange). SH2, Src Homology 2; SH3, Src Homology 3.

There were two mechanisms driving this improved ability of GenomeProt to discriminate expressed proteoforms. Firstly, the RNA-seq-derived database from GenomeProt contains predicted proteins from only expressed transcripts, while UniProt contains isoforms including those from undetected transcripts. For example, the UniProt database contains 8 different AKT2 proteoforms expressed from 10 theoretical *Akt2* transcripts, while the GenomeProt database contains three transcripts identified in the long-read RNA-seq that all encode an identical ORF, allowing peptides to be matched to the only proteoform that is expressed in the sample (**Supplementary Fig. 1g**). Secondly, GenomeProt identifies unique proteotypic peptides from isoforms that were missing in UniProt. In total, we identified 35 peptides from 18 ncORFs encoded by unannotated transcripts not present in GENCODE v35. For example, a novel splice isoform of GPSM2 was validated with an exon-junction-spanning peptide (**Supplementary Fig. 1h**).

To further investigate the potential of using sample-specific long-read RNA-seq-derived GenomeProt databases for proteoform characterisation, we performed an alternative splicing analysis using SUPPA2 (34) which detects 7 types of splicing events: skipping exons (SE), retained intron (RI), mutually exclusive exons (MX), alternative last exons (AL), alternative first exon (AF), alternative 5′ splice site (A5), and alternative 3′ splice site (A3). In total, 40,081 alternative splicing events were identified at the transcript level of which 10,545 had proteomic evidence (**Fig. 2c** and **Supplementary Table 3**). Here, the characterisation of proteoforms is obtained using uniquely mapped peptides that: 1) confirm the existence of a particular exon subjected to splicing, 2) identify unique peptides that span exon-exon junctions, or 3) find unique peptides that map to any downstream exon if the splicing event changes the reading frame and hence translated amino acid sequence. This results in the ability to identify proteomic evidence for either: i) the inclusion transcripts, ii) the exclusion transcripts, or iii) both forms (**Fig. 2c**). Specifically, of the 101,484 uniquely mapping peptides identified in the GenomeProt database, we detected 20,767 peptides spanning exon-exon junctions (**Supplementary Table 2**). AF, SE, A3 and A5 were the most common splicing types at both transcript and protein levels. Both proteoforms were detected at the protein level in approximately 10-25% of the cases. We next used the visualisation module of GenomeProt to provide a deeper exploration of splicing events supported by peptide evidence. Presented examples include two proteoforms of Phospholipid phosphatase 1 (*Plpp1)*generated by mutually exclusive exon use (**Fig. 2d**), two proteoforms from ER Degradation Enhancing Alpha-Mannosidase Like 3 (*E*dem3) generated by alternative A3 splicing (**Fig. 2e**), and two proteoforms of PYM homolog 1, exon junction complex associated factor (*Pym1*) by alternative first exon usage (**Fig. 2f**). We also identified two proteoforms of the Crk-like protein (*Crkl*) generated due to alternative splicing. Domain annotation and structural predictions revealed A0A338P6Q0 lacks the two SH3 domains present in P47941, suggesting these isoforms may have distinct functions (**Figure 2g**). Taken together, these data demonstrate the power of long-read RNA-seq-derived proteogenomics for precise mapping of proteoforms and the utility of GenomeProt for database generation, annotation and isoform visualisation.

### Unannotated peptide detection in the human brain

We next demonstrate the application of GenomeProt to identify unannotated proteins from ncORFs *in vivo*. Here, we examined the human brain, which has a complex and distinct RNA expression and splicing profile (35). We generated paired long-read RNA-seq and proteomic data of three post-mortem human brain regions (frontal cortex [FCX], cerebellum [CBM] and caudate nucleus [CAUD]) from three donors (**Fig. 3a**). GenomeProt identified 89,582 expressed transcript isoforms, including 2,308 isoforms unannotated in GENCODE v47 (total read count across samples >10) (**Supplementary Table S4**). We next used GenomeProt to generate a custom proteome database containing ORFs >30aa, plus uORFs >10aa, resulting in 320,684 entries. Proteomic analysis of each brain region and biological replicate was performed via trypsin/LysC or GluC digestion followed by 2D-LC-MS/MS (3 regions x 3 biological replicates x 2 enzymes x 12 fractions = 216 injections). Data were processed with Spectronaut using the GenomeProt database (see methods). Quality control of the identified transcriptome and proteome confirmed high-quality results with sample separation by brain region based on principal component analysis (PCA) and wide-spread changes in protein abundance (**Supplementary Fig. 2a-b**).

**Figure 3:**
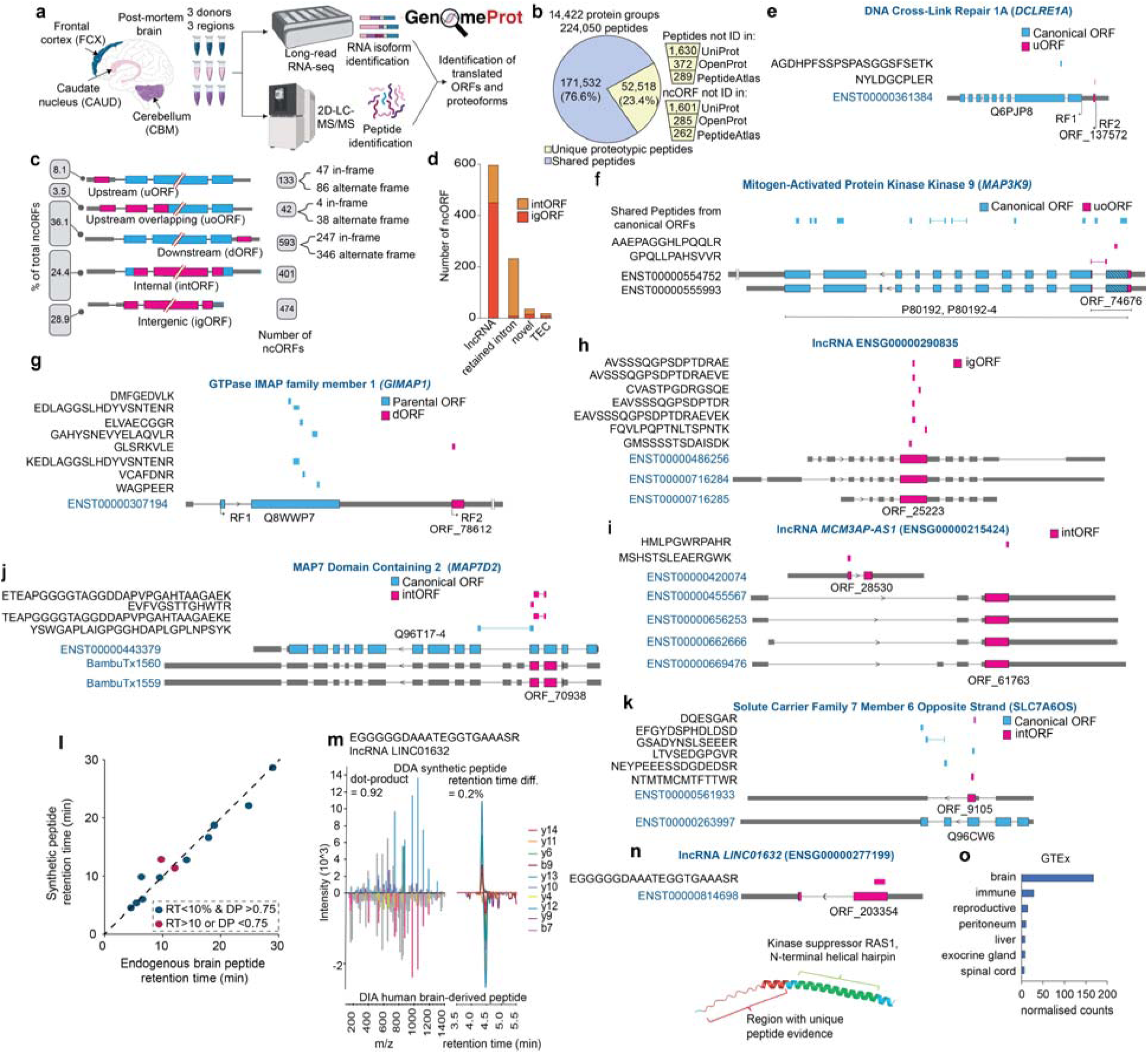
Noncanonical proteins detected with the proteomic evidence in the human brain. (**a**) Experimental design. Expressed transcripts identified based on long-read RNA sequencing from three brain regions — frontal cortex (FCX), caudate nucleus (CAUD), and cerebellum (CBM) from three post-mortem human donors used to construct a customised proteome database, against which peptides identified by 2D-LC-MS/MS were searched. (**b**) Pie chart showing proportion of proteotypic and shared peptides. Inset numbers indicate peptides and non-canonical ORFs (ncORFs) not present in UniProt, OpenProt, and PeptideAtlas databases. (**c**) ncORFs identified based on their genomic and transcriptomic localisation relative to annotated coding regions. uORFs: upstream ORF; uoORF: upstream overlapping ORF; intORF, internal ORF; dORF downstream ORF, and igORF intergenic. Numbers indicate the count of ncORFs in each category; percentages indicate proportion of total ncORFs. (**d**) The number of ncORFs identified across different transcript biotypes, including lncRNA, pseudogene, retained intron, novel, and TEC categories, separated by intORF (bright-orange) and igORF (orange-red). (**e-k**) Representative examples of ncORFs supported by peptide evidence, including a uORF identified upstream of the annotated coding region of DNA Cross-Link Repair 1A (DCLRE1A) (**e**), uoORF identified from alternative frame from Mitogen-Activated Protein Kinase Kinase 9 (MAP3K9), a dORF identified downstream of GTPase IMAP family member 1 (GIMAP1) (**g**), igORF derived from lncRNA ENSG00000290835 (**h**), two intORFs expressed from antisense lncRNA MCM3AP-AS1 (ENSG00000215424) (**i**), an intORF identified from novel transcripts expressed from MAP7 Domain Containing 2 (MAP7D2) (**j**), intORF identified from Solute Carrier Family 7 Member 6 Opposite Strand (SLC7A6OS) (**k**). Parental ORFs and ncORFs are indicated in blue and dark pink, respectively. (**l**) Correlation of peptide spectrum matches (PSMs) between endogenous brain peptides and synthetic peptides. Blue circles indicate peptides with retention time difference <10% and dot-product >0.75; red circles indicate peptides not meeting these criteria. Dashed line indicates perfect correlation. (**m**) Mirror plot comparing MS2 spectra of synthetic peptide (top) and endogenous brain peptide (bottom) from ncORF encoded by *LINC01632*. (**n**) Peptide visualisation on LINC01632-derived ncORF and predicted protein structure using AlphaFold, highlighting the region supported by unique peptide evidence in red and predicted N-terminal helical hairpin domain of kinase suppressor of RAS1 in green. (**o**) GTEx tissue expression profile of lncRNA *LINC01632* across multiple tissues, showing enrichment in brain tissue.

In total, we identified 224,050 peptide sequences on 14,422 protein groups with 52,518 uniquely mapped peptides in brain (**Fig. 3b**). We discovered 2,220 uniquely mapping proteotypic peptides from 1,643 ncORFs which were not annotated in UniProt/RefSeq databases (**Fig. 3b** and **Supplementary Table S5**). Of these, 372 peptides from 285 ncORFs were not mapped to a protein annotated in OpenProt, and 289 peptides from 262 ncORFs were also not identified in the latest Human Peptide Atlas database (Jan 2026) containing >5.3M peptide sequences(36). Furthermore, of the 1,643 ncORFs identified, 9 satisfied the HUPO-HPP criteria for novel protein inclusion (≥2 unique non-overlapping proteotypic peptides each with length of ≥9 amino acids (37), and 37 were reported in a recent large-scale analysis of microproteins and peptideins from the TransCODE Consortium (9). The 1,643 ncORFs included 133 uORFs (47 in-frame and 86 in alternative frames); 42 uoORFs (4 in-frame representing alternative up-stream start sites and 38 in alternative frames); 593 dORFs (247 in-frame and 346 alternate frame); 401 intORFs (excluding ncORFs annotated as predicted isoforms in OpenProt or meeting OpenProt’s criteria for isoforms (38)); and 474 igORFs (**Fig. 3c**). Analysis of transcript biotypes revealed most igORFs represented the translation of previously annotated long noncoding RNAs (lncRNAs). Among the 596 ncORFs encoded by annotated lncRNAs, 450 were igORFs, whereas 146 overlapped annotated protein-coding genes (**Fig. 3d**). We further identified 36 ncORFs from novel transcripts unannotated in GENCODE, including 15 igORFs. We also identified 18 ncORFs from TEC (To Be Experimentally Confirmed; transcripts with protein-coding potential but lacking experimental evidence), of which 8 were intergenic. Additionally, 232 ncORFs originated from transcripts with intron retention, with 224 representing alternative ORFs in annotated protein-coding genes. A total of 17 ncORFs were differentially expressed between the brain regions (**Supplementary Figure 2c**). Examples of ncORFs supported by unique peptide evidence include a uORF identified upstream of the annotated coding region of DNA Cross-Link Repair 1A (*DCLRE1A*) (**Fig. 3e**), uoORF identified from an alternative frame in Mitogen-Activated Protein Kinase Kinase 9 (*MAP3K9*) (**Fig. 3f**), a dORF identified downstream of GTPase IMAP family member 1 (*GIMAP1*) (**Fig. 3g**), an igORF derived from lncRNA ENSG00000290835 (**Fig. 3h**), two intORF expressed from antisense lncRNA *MCM3AP-AS1* (ENSG00000215424) (**Fig. 3i**), an intORF identified from novel transcripts expressed from MAP7 Domain Containing 2 (*MAP7D2*) (**Fig. 3j**), and an intORF identified from Solute Carrier Family 7 Member 6 Opposite Strand (*SLC7A6OS*) (**Fig. 3k**).

We next performed various in silico analyses to investigate the characteristics of ncORFs (**Supplementary Table S6**). The length distribution of ncORFs revealed most of them were <100aa, with median lengths for each category ranging from 33-88aa (**Supplementary Fig. 3a**). Most ncORFs had a higher isoelectric point when compared to those of canonical proteins, suggesting an enrichment of basic, positively charged residues (**Supplementary Fig. 3b**). Hydrophobicity (GRAVY score) was largely comparable between ncORF categories and canonical proteins, with most distributions centred below zero, indicating enrichment of hydrophilic residues. Among ncORFs, dORFs showed a modest shift toward higher hydrophobicity relative to the other categories and to canonical proteins, whereas uoORFs were the most hydrophilic (**Supplementary Fig. 3c**). Consistent with this, ncORFs with lower hydrophobicity tended to have a higher fraction of disordered residues (**Supplementary Fig. 3d**). Subcellular localisation prediction indicated that the majority of ncORFs were predicted to have cytoplasmic and nuclear localisation, with a subset localised to the mitochondrion (**Supplementary Fig. 3e**). To assess ncORF evolutionary conservation, we examined the median ORF phyloP conservation score and the fraction of conserved nucleotides within each ORF. Here, we compared igORFs to canonical proteins across four phylogenetic depths spanning 100 vertebrates, 46 placental mammals, 30 mammals, and 20 primates (**Supplementary Fig. 3f–i**). There was clear separation between canonical and igORFs with few igORFs showing high conservation across vertebrates. In contrast, this distinction was reduced at shallower evolutionary depths. This suggests that igORFs lack the deep evolutionary constraint characteristic of canonical protein-coding regions and are primarily conserved in primates. To further confirm these findings, we performed BLASTP searches comparing igORF-encoded proteins against annotated and predicted (IP_) proteins in the mouse and chimpanzee OpenProt databases. Hits with query coverage and percent identity ≥80% were retained for further analysis. Synteny analysis of these candidates confirmed that 55 igORFs resided within syntenic genomic regions in chimpanzee, whereas none showed significant orthology with mouse, suggesting that these igORFs are primate specific (**Supplementary Table 6**).

To validate the identification of a subset of peptides encoded by lncRNAs, we synthesised 12 peptides and performed LC-MS/MS to compare the retention time and MS/MS fragmentation to the endogenous brain-derived peptide. A total of 10 synthetic peptides eluted with <10% variation in retention time and matched MS/MS fragmentation patterns with a dot-product >0.75 (**Fig. 3l, Supplementary Fig. 4** and **Supplementary Table S7**). For example, a unique peptide mapping to lncRNA *LINC01632* (ENSG00000277199) not previously identified in the Peptide Atlas was validated with a dot-product of 0.92 and retention time <0.2% (**Fig. 3m**). AlphaFold prediction of the protein structure revealed an alpha-helix containing the InterPro homology domain, “Kinase suppressor RAS1, N-terminal helical hairpin”, providing insights into potential functions (**Fig. 3n**). Furthermore, GTEx analysis revealed transcript expression was enriched in the brain (**Fig. 3o**). In summary, paired long-read RNA-seq and deep proteomic data analysed with GenomeProt identified over 1,600 ncORFs in brain demonstrating the discovery power of proteogenomics.

### Identification of missense variants in melanoma treatment resistance

Our third application of GenomeProt focused on the identification of proteoforms created by genetic variants. Both germline and somatic variants can result in amino acid substitutions that alter protein function and may be missed by proteomics using reference databases. In contrast, GenomeProt can identify amino acid changes due to single nucleotide variants (SNVs) from genomic sequencing data and detect their translation into variant peptides and proteins. To demonstrate this application, we investigated the development of treatment resistance in a mouse xenograft model of melanoma. Several resistance mechanisms have been proposed (39), including the presence or acquisition of somatic mutations that allow cancer cell survival and proliferation. However, very few studies have provided protein-level evidence for coding SNVs mediating survival (40), and to our knowledge, none have been validated *in vivo* or in patient samples.

Human *BRAF*^(V600E)^ A375 melanoma cells were subcutaneously implanted into immunocompromised mice and tumours allowed to form prior to treatment with either vehicle or combined Dabrafenib and Trametinib (Dab+Tram) (**Fig. 4a**). Treatment with Dab+Tram initially reduced tumour volume, followed by the development of resistance and tumour regrowth (**Fig. 4b**). Tumours were resected and analysed by whole genome sequencing (WGS), long-read RNA-seq and 2D-LC-MS/MS-based proteomics employing both Trypsin/LysC and GluC digestion. WGS identified 3,797,447 SNVs that passed filters, including 190,361 missense, 9,872 splice-site, and 286,406 UTR variants (**Fig. 4c**). A total of 3,785,127 variants were annotated in the Combined Annotation Dependent Depletion (CADD) database, of which 71,612 had CADD scores ≥ 20, suggesting increased likelihood of disrupting protein expression/function with missense variants having the higher median CADD Phred scores (**Fig. 4d**). To confirm the presence of changes in variant allele frequencies during treatment we compared pre-and early-treatment with the resistance group (**Fig. 4e**). Of the 18,605 variants, 30% showed a >0.1 increase in allele frequency in the resistance group across missense, splice-site and UTR variants, (compared to 5.4% showing a < 0.1 decrease) consistent with the hypothesis of clonal selection and expansion with Dab+Tram treatment (observed as a shift upwards in the y-axis). To perform the GenomeProt analysis, filtered SNVs were inserted into the RNA-seq-derived proteome database using the variant-aware module in GenomeProt and used to process 2D-LC-MS/MS data (see methods). The long-read RNA-seq data identified 116,447 (total read counts across samples > 5, **Supplementary Table S8**) non-redundant transcript isoforms and resulted in a custom proteome database containing 511,621 entries. In total, 78,808 ORFs contained at least one potential SNV. Quality control of the quantified proteome confirmed high-quality results with sample separation by treatment groups based on PCA and the greatest number of protein abundance changes in the Resistance vs Pre-and Early-treatment groups (**Supplementary Fig. 5a-b**). In total, we identified 242,095 peptide sequences on 14,387 protein groups including 5,833 peptides containing SNVs (**Fig. 4f** and **Supplementary Table S9**). We detected the variant BRAF peptide containing the V600E mutation, confirming the validity of the approach (**Fig. 4g-h**).

**Figure 4.**
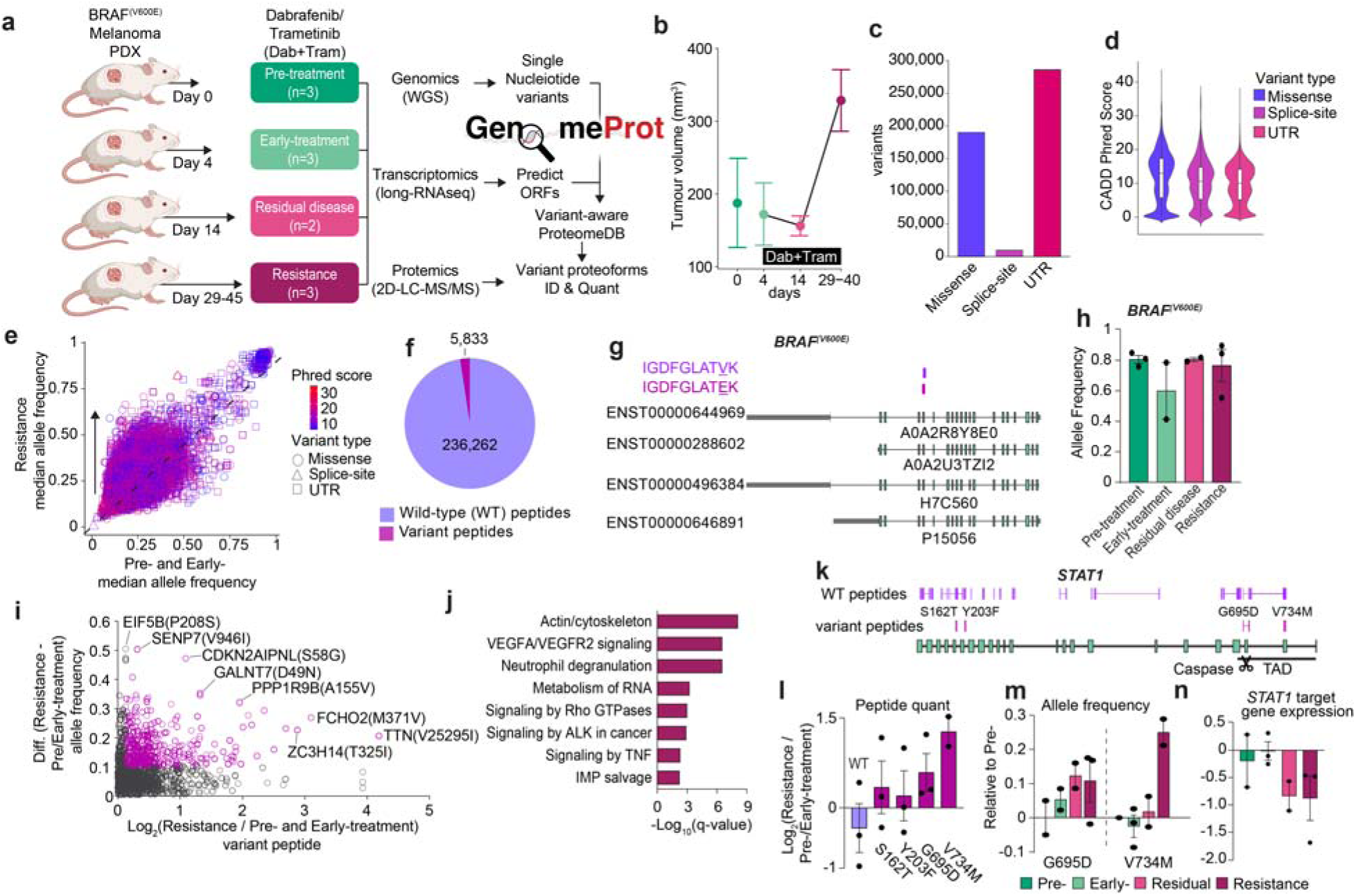
Proteogenomic profiling of *BRAF*^(V600E)^ positive melanoma mouse xenografts and the development of acquired resistance to dabrafenib and trametinib. (**a**) Experimental design. Melanoma mouse xenografts treated with dabrafenib and trametinib (Dab+Tram) were collected at pre-treatment (n = 3), early-treatment (n = 3), residual disease (n = 2) and resistance (n = 3). Variants identified by whole-genome sequencing (WGS) were incorporated into the reference genome, and a variant-aware proteome database was generated from wild-type and variant transcript sequences which was used to search the proteomics data to identify variant peptides. (**b**) Tumour volume measured in pre-treatment, early-treatment, residual disease and resistance groups. (**c-d**). Variant counts and CADD (Phred) score distributions for missense, splice-site and UTR variants included in the variant-aware database. (**e**) Variant allele frequency distribution of missense, splice-site and UTR variants in the resistance group relative to the pre- and early-treatment groups. Circles, triangles, and squares denote missense, splice-site and UTR variants, respectively. Variants are coloured with CADD scores. (**f**) Distribution of variant and wild-type peptides identified by proteomics. (**g**) Wild-type and BRAF^(V600E)^ variant peptides and RNA isoforms. (**h**) Allele frequency of BRAF^(V600E)^ across treatment stages. Bars represent mean values and points indicate individual biological replicates. (**i**) Scatter plot showing differences in variant allele frequency and variant peptide fold change in resistance group relative to pre- and early-treatment groups. (**j**) Top enriched KEGG biological processes linked to genes with variant peptides enriched in the resistance group compared to Pre- and Early-treatment groups. It includes genes with variant peptides showing CADD Phred scores > 10, and a ≥10% increase in allele frequency and variant peptide abundance. (**k**) Identified wild-type (WT) and variant peptides from STAT1. (**l**) Relative abundance of wild-type peptides and peptides harbouring S162T, Y203F, G695D, and V734M variants in the resistance group vs the pre- and early-treatment groups. (**m**) Allele frequencies of the G695D and V734M variants in the *STAT1* gene across Early, Residual disease, and Resistance groups compare to Pre-treatment. (**n**) *STAT1* target gene expression across Pre, Early, Residual disease, and Resistance groups. Error bars indicate standard error of mean.

We next focused on the quantitative analysis of variant peptides by identifying variants with increased allele frequencies during acquisition of resistance to Dab+Tram and also increased in variant peptide abundance hypothesising these may contribute to treatment resistance. A total of 3,569 peptides contained a missense variant that also had an increase in allele frequency in the Resistance group relative to Pre- and Early-treatment groups. We filtered SNVs for a CADD Phred score >10 and a >10% increase in both the allele frequency and variant peptide abundance in the Resistance group and identified 237 peptides from 204 genes, of which 35 have associations to cancer as oncogene or tumour suppressors based on the COSMIC Cancer Gene Census (41), TSGene (42), ONGene (43) or OnkoKB (44) databases (**Fig. 4i** and **Supplementary Table 10**). Pathway enrichment analysis revealed this group was enriched in cellular processes regulating actin/cytoskeleton, neutrophil degranulation, RNA/IMP metabolism and multiple signalling pathways including VEGF, Rho GTPases and ALK, all of which have potential to mediate cellular process for cancer survival (**Fig. 4j**).

Of particular interest were several variants in STAT1, which is a downstream transcriptional effector of all these signalling pathways (**Fig. 4k**). Relative quantification of wild-type STAT1 protein levels with ∼50 non-variant peptides suggested a potential subtle decrease in the Resistance group relative to Pre- and Early-treatment (Log_2_[fold-change] = −0.34) (**Fig. 4l**). In contrast, we identified four *STAT1* peptide variants, including G695D and V734M in the C-terminal transcriptional activation domain (TAD) that increased in the Resistance group by >50%. Consistent with the peptide findings, the allele frequencies of both variants also increased throughout development of resistance (**Fig. 4m**). Both variants are particularly interesting as D694/G695 is a validated Caspase cleavage site that regulates *STAT1* activity (45), and V734 is directly adjacent to a phosphorylation hotspot required for the maximal activation of *STAT1* [PMID: 7543024], and hence mutation may impair activity. Indeed, enrichment analysis in the long-read RNA-seq data revealed that *STAT1* target gene expression was decreased in the Resistance group suggesting reduced *STAT1* activity consistent with a previously described role as a tumour suppressor and potential role in immune evasion (**Fig. 4n**) (46, 47). Taken together, we demonstrate the utility of the variant-aware module of GenomeProt to incorporate variants identified by WGS into the proteogenomics analysis and identify peptide/protein variants that have likely roles in the acquired resistance to melanoma treatment.

## DISCUSSION

Proteogenomics is a powerful technique for transcriptome and proteome investigation, but requiring knowledge across transcriptomics, proteomics and bioinformatics can make it difficult to perform. We present GenomeProt, a GUI-based software package designed to make proteogenomics broadly assessable to researchers. GenomeProt includes features to carry out all the key steps in proteogenomics including generating an *in silico*, sample-specific, proteome database derived from RNA-seq; peptide mapping to ORFs; proteogenomic data integration; and uniquely, the ability to perform genome- and transcriptome-centric visualisation of identified peptides, ORFs and proteins. Our visualisation module allows easy searching and filtering for genes of interest to examine peptide-level data on: i) canonical proteoforms, ii) ncORFs, and iii) amino acid variants. GenomeProt was designed so that users with reasonable proteomic experience, but a limited genomics/transcriptomics background and vice-versa could perform proteogenomics, while at the same time not requiring advanced bioinformatics analysis skills. We achieved this via a GUI-based workflow and the ability to work with multiple datatypes (FASTQ, BAM or GTF). GenomeProt produces multiple data outputs including a FASTA for mass spectrometry analysis, which includes sequence header annotations, and metadata containing additional ORF annotations for downstream analysis. The proteogenomic data integration and visualisation modes of GenomeProt produce a seamless analysis suite for genome mapping and interrogation of specific proteoform translation and relative abundance.

GenomeProt enabled comprehensive proteoform characterisation. Benchmarking GenomeProt via a proteogenomic analysis of Kusa 4B10 cells revealed a similar number of peptides and proteins were identified compared to canonical UniProt/OpenProt protein databases. However, GenomeProt identified more unique proteotypic peptides enabling superior discrimination and precise identification of proteoforms. Furthermore, annotation of >20,000 exon-junction-spanning peptides within GenomeProt, combined with an analysis of alternative splicing, identified protein-level evidence for >10,000 alternative splicing events. Equivalent analyses would be particularly powerful for the characterisation of isoform switching events following changes in alternative splicing during normal healthy development or during the acquisition of disease. Furthermore, it would be interesting to apply GenomeProt to further understand the mechanisms regulating alternative splicing by manipulating the expression of splicing factors and then performing proteogenomic analysis. Finally, the integration of machine learning models that predict functional outcomes of splicing (48) with experimental data generated with GenomeProt could enhance the development of isoform-switching therapeutics.

The international TransCODE consortium (49) was established to set up a pathway and standards for inclusion of uncharacterised proteins from ncORFs into formal reference gene annotations. This is important given the large diversity of identification and annotation criteria in recent community evaluation studies(50) and the thousands of ncORFs identified by TransCODE itself (9). Motivated by the TransCODE Consortium (49), one of our goals was to increase routine detection of ncORFs by facilitating the ease of proteogenomic analysis. We demonstrated this by using GenomeProt to identify ncORFs in human brain. We identified hundreds of potential ncORFs and validated ten that were expressed from previously annotated lncRNAs. For example, we validated a ncORF expressing lncRNA *LINC01632* (ENSG00000277199) that is enriched in the brain and contains a Kinase suppressor RAS1, N-terminal helical hairpin domain that shares sequence similarity with the KSR1 gene. This domain is known to dimerise with SAM-like domains (51) and mice lacking *Ksr1* (MGI Ref. ID: J:75739) have impaired synaptic plasticity and reduced long-term potentiation (52). Despite the high validation rate with synthetic peptides, it is important that the additional ncORFs identified in the current study are further analytically validated. Furthermore, annotation of these as protein-coding genes has historically required the validation of biological function. Hence, the classification of analytically validated ncORFs as peptideins as an intermediatory annotation is warranted: confidently detected but their biological function is currently unknown. One important step in assigning function is routine quantification to observe changes in abundance during healthy development or disease. Hence, GenomeProt represents a tool to perform such routine quantification across laboratories.

In this study we focused on transcriptomic analysis using long-read RNA-seq that provides advantages for proteoform annotation. However, proteogenomics using transcriptomic data generated by short-read Ribo-seq has the advantage of capturing actively translated transcripts for ncORF analysis. Here, ribosome-protected RNA fragments or ‘footprints’ are purified to identify ORFs present on translating ribosomes. However, the short length of Ribo-seq reads is a limitation for identifying the translated ORFs and proteoforms and Ribo-seq data has an inherent 5′ bias in read coverage, which may skew uniformity (3). Future work building *in silico* predicted protein databases by integrating long-read RNA-seq with Ribo-seq from the same sample(s) are warranted and will undoubtably improve the quality of proteogenomics. Furthermore, we demonstrated the utility of GenomeProt with protease-treated ‘bottom-up’ proteomic analysis. Integration of these data with human leukocyte antigen (HLA)-enriched immunopeptidomics(53, 54) or ‘middle-down’ proteomics (55), are also likely to improve both coverage and confidence of ncORF analysis.

One unique feature of GenomeProt compared to existing tools is the ability to include variants via VCF input. We applied this workflow to identify and quantify changes in the abundance of variant peptides during the development of melanoma treatment resistance. We identified variant peptides from 35 tumour associated genes (oncogenes or tumour suppressors) that increased in abundance with resistance, and the SNV encoding these amino acid changes also increased in allele frequency during the treatment. Of particular interest were two variants in the C-terminal transcriptional activation domain (TAD) of STAT1 that increased during resistance and were associated with a decrease in STAT1 target gene mRNA expression, which is suggestive of decreased STAT1 activity. STAT1 has a complex role in cancer biology, including as a tumour suppressor, and elevated expression is generally associated with improved prognosis in several cancers (56, 57). The mechanisms are complex, but the potential decreased activity observed during the development of resistance may facilitate escape from cell death pathways or immune surveillance, promoting cellular proliferation and survival.

While GenomeProt enables variant aware proteogenomics, there are limitations in the type and complexity of genomics variants it can incorporate. Future iterations could include the phasing of variants and optionally allow multiple variant alleles. This could be beneficial as we observed multiple variants in 6 of the 35 tumour associated genes. While melanoma has one of the highest SNV rates (58), multiple somatic mutations in the same ORF and within the same cell are potentially rare. If an ORF has been identified with multiple somatic mutations via, e.g., bulk tissue WGS, it is more likely that these arise in different cell populations and will generate different variant-containing proteoforms. However, GenomeProt currently includes all identified variants in an ORF as a single entry in the database, i.e.: all ORFs containing multiple variants are represented by two entries; wild-type and the in silico predicted sequence containing all identified variants. Hence, this likely underestimates the complexity of potential variant-containing proteoforms present. We accepted this trade-off as including each missense variant as a separate entry combined with three-frame translation would result in an unreasonable increase in database size and cause a decrease in peptide mapping sensitivity due to the increased cutoffs required to achieve desired FDRs.

GenomeProt greatly increases the accessibility of proteogenomics with its local GUI and public browser versions. However, a limitation of our hosted browser version is it doesn’t accept RNA-seq FASTQs as input or perform the proteomics analysis step due to the size of FASTQ files and the computationally intensive nature of mapping RNA-seq reads and performing the proteomics step. Hence, generation of BAM files is required prior to data upload and an external proteomics pipeline must be used. Alternatively, local installation avoids these limitations as users can use FASTQ as data input and use the FragPipe proteomics pipeline integrated into GenomeProt. For analysis of multiple biological samples, we chose to generate a single FASTA file containing aggregated *in silico*-predicted ORFs from all long-read RNA-seq runs. This means all mass spectrometry files from the various samples can be analysed in a single batch against this aggregated database. For studies involving multiple biological samples, users have the ability to generate individual FASTA files that are sample-specific. However, this would require separate mass spectrometry searches for individual samples and therefore, careful consideration of how the mass spectrometry data are aggregated and post-processed for e.g. accurate quantification and FDR calculation(59). This is particularly important for proteogenomic studies focused on personalised/precision medicine or analysis of rare diseases from individual patients.

In summary, we present GenomeProt, which aims to overcome the barriers to routine use of proteogenomics to more accurately characterise the proteome. We compare its performance against standard canonical UniProt/OpenProt and show improved proteoform characterisation and utility in the analysis of alternative splicing. We also demonstrate GenomeProt can identify ncORFs including novel peptides in the human brain, a subset of which we validate by analysis of synthetic peptides. The inbuilt visualisation module of GenomeProt is a unique feature that allows peptide mapping onto the genome and transcriptome for detailed characterisation of proteoforms. Finally, inclusion of amino acid variants into FASTA files via direct VCF incorporation adds an additional unique functionality of GenomeProt and we demonstrate its utility by performing a proteogenomic analysis of melanoma treatment resistance.

## METHODS

### Implementation of GenomeProt code

GenomeProt provides a user-friendly graphical user interface (GUI) implemented in R Shiny and includes an installer script that automatically configures a Conda environment with all required dependencies. The complete GenomeProt application is also distributed as a docker container. The database generation module accepts multiple input formats, including raw FASTQ files, pre-aligned BAM files, or user-supplied GTF files with transcript models derived from the samples. GenomeProt supports both short-read and long-read RNA-seq data. For short-read FASTQ inputs, the tool generates a custom index for the user-specified genome and transcriptome, followed by alignment and quantification using Salmon’s selective-alignment approach. Users can provide transcriptome-aligned BAM files generated by STAR (60) or HISAT2 (61) in the web server, which are subsequently quantified using Salmon (v1.10.3) (23). For long-read RNA-seq data, GenomeProt aligns FASTQ files using Minimap2 (v2.28) (24), followed by transcript identification and quantification using bambu with default parameters, which automatically estimates a dataset-specific Novel Discovery Rate (NDR) for transcript discovery (v3.4.0) (25). Bambu can also identify novel/unannotated transcripts in addition to annotated transcripts. Low-abundance transcripts are filtered using a user-defined minimum total read count across all samples (default: 5). Transcripts passing this threshold undergo in silico translation in all three reading frames using ORFik (v 1.22.0) (**62**). Resulting ORFs are clustered with CD-HIT (v4.8.1) (63), retaining the longest representative sequence. ORFs from alternative reading frames of annotated coding (CDS) regions in mRNAs are excluded. The database generation module produces a multi-FASTA database file along with a metadata file with additional information on annotated proteins (including if the protein/RNA is present in UniProt, RefSeq, or GENCODE) and unannotated proteins, including OpenProt identifiers for proteins detected in the OpenProt database. GenomeProt enables variant-aware proteome database generation. Homozygous and heterozygous variants from a user-supplied combined VCF file are separated and incorporated into the reference genome using bcftools consensus (v1.21)(64), generating two customised genome FASTA: one containing only homozygous variants and another containing both homozygous and heterozygous variants. Corresponding transcriptome databases with wildtype and variant transcripts are generated which are then used to produce wild-type and variant ORFomes, which undergo the same downstream processing as the standard GenomeProt proteogenomics database. The proteomics analysis module runs FragPipe (29) and requires users to input basic mass spectrometry parameters including the proteases used. The integration module processes peptide-level data from proteomic search outputs, maps peptide coordinates onto spliced transcript coordinates, and generates a GTF file containing genomic annotations for both peptides and transcripts. The visualisation module leverages a custom version of IsoVis (65), a website written in Nuxt 2 and originally developed for visualising alternative RNA isoforms, to enable the interactive exploration of peptide mappings and associated transcript features. The combined GTF file with peptide and transcript coordinates outputted by the integration module, and optionally the transcript counts used for the database generation module and the peptide intensities from proteomics searches, are provided as inputs. Users can query a gene of interest by providing a gene symbol or Ensembl gene ID and by applying various filters to restrict the analysis to specific genes, such as those with uniquely mapped peptides. Transcripts, ORFs and peptides are represented as separate tracks. Users can optionally visualise transcript and peptide abundance across samples.Visualisations can be customised by toggling the visibility of certain plots, enabling intron normalisation to display introns with uniform widths, and switching the reading direction. Users can zoom into plots by selecting a region or specifying a coordinate range. Isoforms, ORFs, and peptides can be reordered, sorted alphabetically, and toggled on or off. Isoforms and peptides can also be sorted by their heatmap mean values. Visualisations can be exported as PDFs, SVGs, PNGs, and JPEGs to create publication-ready images.

### RNA isolation from Kusa 4B10 cells

Female Kusa 4B10 stromal stem cells were grown in Minimum Essential Medium alpha (MEMalpha) (GIBCO by Life Technologies; #12561056) supplemented with 10% fetal bovine serum (FBS) (Life Technologies; #26140079), pyruvate and GlutaMAX (GIBCO by Life Technologies). Cells were kept at 37°C and 5% CO2 in a humidified incubator Direct Heat CO2 Incubator featuring Oxygen Control (In Vitro Technologies). All cell stocks were regularly checked for absence of mycoplasma with the Mycoplasma Detection Kit (Jena Bioscience; # PP-401). Cells were washed twice with cold PBS to remove culture media and immediately lysed in 500 μl cold QIAzol reagent with additional agitation using a disposable 1.5 mL tube pestle. RNA was isolated from lysate using the RNeasy Lipid Tissue Kit (QIAGEN, 74804) following the manufacturers’ instructions with minor changes. Briefly, 100 μl Chloroform - isoamyl alcohol (Merck, 25666) was added to each lysed sample, and the resultant upper aqueous phase was collected. 70% molecular grade ethanol was added 1:1 to the aqueous phase collected. RNA was eluted in 40 μl nuclease-free water. The purified RNA quality and quantity was checked using a Qubit 4 RNA BR (Thermo Fisher Scientific, Q10211) kit (1 μL), RNA ScreenTapes (1:4) (Agilent, 5067-5576) on a TapeStation 4200 (RINe cut-off 8) and Nanodrop 2000 (1 μL).

### RNA isolation from human post-mortem brain

Nine control post-mortem human brain samples were obtained from five individuals collected through the Victorian Brain Bank (VBB) under HREC approvals #12457 and #28304. Briefly, samples comprised 4 males and 1 female, age range: 64 – 81.2 yrs, PMI range: 24 – 64 hrs. Frozen tissue (weight range: 37 – 102 mg) was cut from three brain regions (N=3 each) including BA46 (medial frontal cortex), caudate nucleus and cerebellum. Total RNA was extracted from bulk tissue in batches of 3 – 6 samples. First, frozen brain tissue was homogenised on ice, using a manual tissue grinder (Potter-Elvehjem, PTFE), whilst immersed in 1 mL QIAzol Lysis Reagent. Lysate was then processed using a RNeasy Lipid Tissue Kit (QIAGEN, 74804), according to the manufacturers’ instructions. Isolated RNA quality and quantity was checked using a Qubit 4 RNA BR (Thermo Fisher Scientific, Q10211) kit (2 μL), RNA ScreenTapes (Agilent, 5067-5576) on a TapeStation 4200 (RINe cut-off = 6) and Nanodrop 2000.

### RNA and genomic DNA (gDNA) isolation from a mouse melanoma xenograft model

Melanoma xenograft experiments were performed according to protocols approved by the Animal Ethics Committee of Peter MacCallum Cancer Centre and in accordance with the National Health and Medical Research Council Australian code for the care and use of animals for scientific purposes, 8^th^ Edition, 2013. Human *BRAF* mutant melanoma A375 cells were prepared in 50% Matrigel in PBS and 4 million cells were subcutaneously injected into the right flank of 6–8 week-old male BalbC nu/nu (nude) mice. Once tumours reached a mean volume of 150-200mm^3^, mice were randomised into 5 groups (N=4) and were treated with either vehicle (0.5% HPMC and 0.2% Tween 80 in H20) or Dabrafenib (Dab; 30mg/kg in 0.5% HPMC and 0.2% Tween 80 in H20) and trametinib (Tram;0.15mg/kg in 0.5% HPMC and 0.2% Tween 80 in H20), administered by oral gavage daily for 6 out of 7 days each week. Mice were euthanised and tumours harvested at pre-determined timepoints; pre-treatment (0 days), early-treatment (vehicle or Dab/Tram for 4 days), residual disease (Dab/Tram for 14 days) and resistance (defined as tumour regrowth under therapeutic pressure; mean volume ∼200-400mm3). Vehicle controls for residual disease and resistant groups were not included due to ethical endpoints for maximum tumour size (1200mm3). Tumours were cut into pieces (where possible), snap frozen and stored at −80oC. Prior to isolation of RNA or gDNA, tumour pieces were crushed on dry ice and lysed using kit-specific lysis buffers and homogenised using an Eppendorf pestle. RNA isolation was performed using the MACHEREY-NAGEL NucleoSpin RNA kit (#740955.250) as per manufacturer’s directions. RNA quantity was determined using a Qubit 4 RNA BR (Thermo Fisher Scientific, Q10211) kit, and quality was assessed using RNA ScreenTapes (Agilent, 5067-5576) on a TapeStation 4200 (RINe cut-off = 9). gDNA was isolated using a DNeasy Blood & Tissue Kit (QIAGEN, 69504) as per manufacturer’s directions. gDNA quantity was determined using a Qubit dsDNA BR Assay (Thermo Fisher Scientific, Q33266) and quality was assessed using gDNA ScreenTapes (Agilent Technologies, 5067-5365) on a TapeStation 4200 (Agilent).

### Kusa 4B10 cell Nanopore whole transcriptome (cDNA-PCR) library preparation and sequencing

500 ng of Kusa total RNA for each sample was used for Nanopore long-read library preparation which followed the recommended cDNA-PCR (Oxford Nanopore Technologies, SQK-PCB114-24) sequencing protocol. Barcoding PCRs were done in duplicate using LongAmp Hot Start Taq 2X Master Mix (NEB, M0533) with a 7 min extension time (approximately 1 min/Kb) and 12x cycles. Cleaned PCR was checked for yield and length profile using Qubit 1X dsDNA HS Assay Kit (Invitrogen, Q33231) and Genomic DNA ScreenTapes (Agilent Technologies, 5067-5365), respectively. Three randomised Kusa samples were prepared and pooled equally (∼16.6 fmol). Rapid adaptor ligated libraries were loaded (∼50 fmol) onto a PromethION flow cell (FLO-PRO114M) and run for 72 hrs. All runs were basecalled live (MinKNOW v24.06.10) using the high-accuracy (HAC v4.3.0, 400 bps) basecalling model (Dorado v7.4.12) and minimum pass qscore = 9.

### Human brain Nanopore whole transcriptome (cDNA-PCR) sequencing

500 ng of PM brain total RNA for each sample was used for Nanopore long-read library preparation which followed the recommended cDNA-PCR (Oxford Nanopore Technologies, SQK-PCB114-24) sequencing protocol with minor changes. Each sample included a Spike-In RNA Variant (SIRV) mix from SIRV-Set 1 (Lexogen, 025.03) to monitor sensitivity and aid detection and quantification of isoform expression between sample types. Mix E1 was spiked into frontal cortex and caudate samples and Mix E2 was spiked into cerebellum samples. Barcoding PCRs were done in duplicate using LongAmp Hot Start Taq 2X Master Mix (NEB, M0533) with a 7 min extension time (approximately 1 min/Kb) and 12x cycles. Cleaned PCR was checked for yield and length profile using Qubit 1X dsDNA HS Assay Kit (Invitrogen, Q33231) and Genomic DNA ScreenTapes (Agilent Technologies, 5067-5365), respectively. Three randomised brain samples were prepared and pooled equally. Rapid adaptor ligated libraries were loaded (∼50 fmol) onto a PromethION flow cell (FLO-PRO114M) and run for 72 hrs. All runs were basecalled live (MinKNOW v24.06.16) using the super-accurate (SUP v4.3.0, 400 bps) basecalling model (Dorado v7.4.14) and minimum pass qscore = 10.

### Nanopore whole transcriptome (cDNA-PCR) sequencing for melanoma tumours

300 ng of RNA was used for library preparation according to the PCR-cDNA Barcoding Kit (Oxford Nanopore Technologies, SQK-PCB111.24). Briefly, cDNA was prepared and stored overnight at 4°C, prior to PCR amplification using LongAmp Hot Start Taq 2X Master Mix (NEB, M0533) with a 7 min extension time and 14x cycles. Purified PCR was checked for yield and length profile using Qubit 1X dsDNA HS Assay Kit (Invitrogen, Q33231) and Genomic DNA ScreenTapes (Agilent Technologies, 5067-5365), respectively. Two randomised samples were pooled (1:1) to a total of 30 fmol and loaded onto a PromethION flowcell and sequenced on a P2 Solo Oxford Nanopore platform (PRO-SEQ002) for up to 72hrs. All runs were basecalled live (MinKNOW v24.06.16) using the super-accurate (SUP v4.3.0, 400 bps) basecalling model (Dorado v7.4.14) and minimum pass qscore = 10, producing on average 40 million passed reads per sample. Data were analysed following the nfcore-nanoseq workflow (https://github.com/nf-core/nanoseq), with the addition of xenomapper (<u>xenomapper.nf</u>) to explore the proportion of reads arising from human melanoma cells versus the mouse host (https://github.com/Oshlack/xenanoseq). For the analysis reported here, we have used bam files generated from all reads. Alignment was performed using minimap2 (v2.17) and transcript discovery and quantification was performed using bambu (v3.4.0) and the human genome (GRCh38) as a reference.

### Illumina whole genome sequencing for melanoma tumours

Whole genome sequencing was performed by the Molecular Genomics Core Facility at the Peter MacCallum Cancer Centre and the Australian Genomics Research Facility (AGRF). Briefly, 1µg gDNA was used for library preparation according to the NEB DNA Ultra II kit and was sequenced at ∼30x coverage using 3 lanes of a NovaSeq X 10B flowcell.

### Proteomic sample preparation from Kusa cells

Kusa 4B10 stromal stem cells (SSCs) were differentiated into osteoblasts over 21 days. Cells were washed twice with cold PBS and scraped in 4% sodium deoxycholate (SDC) in 100 mM Tris pH 8.5 and tip-probe sonicated. Proteins were quantified with BCA protein assay kit (ThermoFisher Scientific), reduced with 10 mM Tris(2-carboxyethyl)phosphine hydrochloride (TCEP) and 40 mM 2-Chloroacetamide (CAA) for 5 min at 95 degrees. The samples were cooled and digested with sequencing-grade trypsin (Promega) and Lys-C (Wako, Japan) or sequencing grade GluC or AspN (Promega) (protein:enzyme ratio 100:1) for 16 h at 37 °C with shaking at 1600 rpm. Peptides were purified using in-house Styrenedivinylbenzene-Reversed Phase Sulfonate (SDP-RPS) stage tips and eluted with v/v 80% acetonitrile/5% NH4OH at 500 g for 5 min. Eluted peptides were dried in a vacuum concentrator (Eppendorf, Germany) at 45 °C for 45 min and the dried peptides were resuspended in 2% acetonitrile/0.1% TFA. Peptides were fractionated by neutral phase C18BEH HPLC into 12 fractions as previously described (66).

### Proteomic sample preparation of human brain

Protein extractions were carried out on the same brain regions from the same individual donors as utilised for RNA-seq. Frozen brain samples were lysed in 6 M guanidine HCl (Sigma), 100 mM Tris pH 8.5 containing 10 mM TCEP and 40 mM 2-chloroacetamide (CAA)by tip-probe sonication. The lysate was heated at 95 °C for 5 min and centrifuged at 20,000 x g for 10 min at 4 °C. The supernatant was diluted 1:1 with water and precipitated overnight with 5 volumes of acetone at –20 °C. The lysate was centrifuged at 12,000 x g for 5 min at 4 °C and the protein pellet was washed with 80% acetone. The lysate was centrifuged at 12,000 x g for 5 min at 4 °C and the protein pellet was resuspended in Digestion Buffer (10% 2,2,2-Trifluoroethanol (Sigma) in 100 mM HEPEs pH 8.5). Protein was quantified with BCA (ThermoFisher Scientific) and digested with either sequencing-grade trypsin (Sigma) and Lys-C (Wako) or GluC (Promega) at a 1:50 enzyme:substrate ratio overnight at 37 °C with shaking at 2000 x rpm. Peptides were purified using in-house Styrenedivinylbenzene-Reversed Phase Sulfonate (SDP-RPS) stage tips and eluted with v/v 80% acetonitrile/5% NH4OH at 500 g for 5 min. Eluted peptides were dried in a vacuum concentrator (Eppendorf, Germany) at 45 °C for 45 min and the dried peptides were resuspended in 2% acetonitrile/0.1% TFA. Peptides were fractionated by neutral phase C18BEH HPLC into 12 fractions as previously described (66).

### Proteomic sample preparation of melanoma tumours

Frozen melanoma samples were lysed in 6 M guanidine HCl (Sigma), 100 mM Tris pH 8.5 containing 10 mM TCEP and 40 mM 2-chloroacetamide (CAA)by tip-probe sonication. The lysate was heated at 95 °C for 5 min and centrifuged at 20,000 x g for 10 min at 4 °C. The supernatant was diluted 1:1 with water and precipitated overnight with 5 volumes of acetone at –20 °C. The lysate was centrifuged at 12,000 x g for 5 min at 4 °C and the protein pellet was washed with 80% acetone. The lysate was centrifuged at 12,000 x g for 5 min at 4 °C and the protein pellet was resuspended in Digestion Buffer (10% 2,2,2-Trifluoroethanol (Sigma) in 100 mM HEPEs pH 8.5). Protein was quantified with BCA (ThermoFisher Scientific) and digested with either sequencing-grade trypsin (Sigma) and Lys-C (Wako) or GluC (Promega) at a 1:50 enzyme:substrate ratio overnight at 37 °C with shaking at 2000 x rpm. Peptides were purified using in-house Styrenedivinylbenzene-Reversed Phase Sulfonate (SDP-RPS) stage tips and eluted with v/v 80% acetonitrile/5% NH_4_OH at 500 g for 5 min. Eluted peptides were dried in a vacuum concentrator (Eppendorf, Germany) at 45 °C for 45 min and the dried peptides were resuspended in 2% acetonitrile/0.1% TFA. Peptides were fractionated by neutral phase C18BEH HPLC into 12 fractions as previously described (66)

### Mass spectrometry-based proteomics and data processing

Peptides were analysed on Vanquish Neo nanoUHPLC coupled to an Orbitrap Astral mass spectrometer (ThermoFischer Scientific) via electrospray ionisation in positive mode with 1.9 kV at 275 °C and RF set to 50%. Separation was achieved on a 5.5 cm uPAC Neo column (ThermoFischer Scientific) over 23 min at a flow rate of 750 nL/min. The peptides were eluted over a linear gradient of 3–23% Buffer B (Buffer A: 0.1% v/v formic acid; Buffer B: 90% v/v acetonitrile, 0.1% v/v FA) and the column was maintained at 50 °C. The instrument was operated in data-independent acquisition (DIA) mode with an MS1 spectrum acquired over the mass range 380-980 *m/z* (120,000 resolution, 500% automatic gain control (AGC) and 5 ms maximum injection time) followed by MS/MS with 300 x 2 *m/z* isolation (500% AGC, 27% normalised collision energy (NCE) and 3 ms injection time) and a loop time of 0.6 s. Synthetic peptides were analysed with identical HPLC method and MS acquisition settings except the data were acquired with data-dependent acquisition. Raw DIA data were processed with Spectronaut (v20.3.251215) with default parameters with peptide spectral match FDR, peptide FDR and protein group FDR set to 0.01. Dynamic MS1 and MS2 mass tolerance was enabled, and retention time calibration was accomplished using local (non-linear) regression. The default dynamic extracted ion chromatogram window size was performed. Peptide quantification was carried out at MS2 level using 3-6 fragment ions, with automatic interference fragment ion removal as previously described (67). The raw data for each protease were searched separately and results combined to create communal library and repeat protein inference analysis using “From search engine” setting and re-calculation of protein group-level FDR. Peptide reports were exported and uploaded to GenomeProt for genome annotation and visualisation. DDA data from synthetic peptides were processed using Skyline (v21.1.0.146) (68) and spectral libraries built with MSFragger within FragPipe (v18) (29) using BiblioSpec (69). Precursor and product ion extraction ion chromatograms (XICs) were generated using extraction windows two-fold the full width at half-maximum for both MS1 and MS2 filtering. Ion match tolerance was set to 0.055 m/z and matched to charges 2+ and 3+ for MS1 filtering of the first three isotopic peaks and 1+ and 2+ for MS2 filtering of b-and y-type ions. All data were manually confirmed for co-elution of MS1 and MS2 and have been uploaded to the Panorama Repository (see Data and Code Availability section).

### Custom proteogenomic database generation

#### Mouse proteogenomics database

BAM files generated from SUP basecalling with Dorado (v0.7.0) were used to construct a custom proteome database using GenomeProt, with GENCODE (v35) as the reference annotations. The database was generated using a minimum transcript expression threshold >5, ORF length >30 amino acids, and inclusion of short uORFs >10 amino acids.

#### Human brain proteogenomic database

BAM files generated with Dorado (v0.7.0) were used to build a custom proteome database, with GENCODE (v47) as the reference annotations. Parameters were the same as for the mouse dataset, except that the minimum transcript expression threshold was set to >10.

### Melanoma variant-aware proteogenomics database, variant calls generation and processing

Reads in the FASTQ files generated from whole-genome sequencing (WGS) were aligned to the human reference genome (GRCh38) using BWA-MEM2 (v2.2.1) (70). Adapter sequences were trimmed using Fastp (v0.23.2) (71), and quality control was performed using FastQC (v0.12.1). The resulting BAM files were sorted using SAMtools (v1.16.1) (72) and pre-processed according to the GATK Best Practices workflow (v4.3.0.0) (73), including duplicate marking using MarkDuplicates, base quality score recalibration (BQSR), and BAM indexing. Variant calling was performed using Mutect2. Individual VCF files were merged using bcftools merge (64), resulting in a merged VCF file containing 10,809,810 variants. The merged VCF file was further annotated using VEP (v107) (74). Variants detected in at least two samples within any group were retained for downstream analysis, resulting in 6,006,236 SNVs. Subsequently, synonymous, stop, intronic, and NMD transcript variants were filtered, resulting 3,797,447 SNVs that were used to generate variant-aware database.

### Downstream analysis

#### Splicing analysis of Kusa data

Alternative splicing events were identified using SUPPA2 (v2.3) from a GTF file of expressed transcripts generated by bambu (v3.4.0) using Kusa long-read RNA-seq data. Novel transcripts (BambuTx accessions) were defined relative to the GENCODE (v35) reference transcript annotations. The generateEvents module of SUPPA2 was used to identify skipped exons (SE), mutually exclusive exons (MX), alternative 5′ splice sites (A5), alternative 3′ splice sites (A3), retained introns (RI), alternative first exons (AF), and alternative last exons (AL), which were exported in IOE format for downstream analyses. Integration of splicing and peptide data was performed using a custom python script.

#### Sequence feature analysis of ncORF-derived proteins from human brain proteogenomics data

Subcellular localisation of ncORF-derived proteins across different categories, including uORFs, uoORFs, dORFs, intORFs, and igORFs, was predicted using DeepLoc 2.1 (75) Intrinsically disordered regions (IDRs) were identified using IUPred2A (76). Conserved protein domains in igORF-derived proteins were predicted using InterProScan (v6.0.1) (77).

#### Conservation analysis of igORFs and canonical coding regions

PhyloP conservation tracks in bigWig format (.bw) were obtained from the UCSC Genome Browser (78), including 100 vertebrates, 46 placental mammals, 30 mammals, and 20 primate conservation datasets(79, 80). Conservation analysis was restricted to igORFs, as other ncORF categories may overlap with annotated coding regions, where conservation signals could be influenced by existing protein-coding sequences. Genomic coordinates for canonical ORFs and igORFs were obtained from the proteome database metadata file. All canonical ORFs were included in the analysis irrespective of peptide support from proteomics datasets. The median PhyloP score was calculated for igORFs and canonical ORFs. The fraction of conserved nucleotides was calculated as the proportion of nucleotides with PhyloP scores exceeding predefined thresholds (0.2 for 100 vertebrates and 0 for 46 placental mammals, 30 mammals, and 20 primates) relative to the total number of nucleotides in the ORF. The distribution of per-ORF median PhyloP conservation scores and the fraction of conserved nucleotides within ORFs were compared across different conservation datasets.

#### Protein orthology and synteny analysis of igORFs

igORF protein sequences were compared against annotated and predicted (IP_) proteins in the mouse (Mus musculus) and chimpanzee (Pan troglodytes). Custom blast databases were built from OpenProt databases with the makeblastdb utility from standalone BLAST+ (v2.17.0) (81). igORF sequences were queried against these databases using BLASTP with the following parameters: -word_size 3 -matrix BLOSUM62 -gapopen 11 -gapextend 1 -evalue 1e-5 -max_target_seqs 5. The best hit for each igORF with query coverage and percent identity ≥80% was retained for downstream synteny analysis.

To determine whether igORFs occupied syntenic genomic loci in chimpanzee, we used the UCSC hg38ToPanTro5 chain file, which stores the coordinates of aligned human-chimpanzee genomic blocks. Alignment blocks corresponding to the genomic coordinates of each igORF (hg38) were retrieved using the liftOver function in the rtracklayer R/Bioconductor package. Genomic coordinates of the orthologous chimpanzee protein identified from the BLASTP analysis were then compared against these extracted alignment blocks. A locus was classified as syntenic if the chimpanzee ORF coordinates were either fully contained, or overlapped with, at least one extracted alignment block.

#### Differential expression analysis of proteins expressed in human brain regions

Quantification data were obtained by searching proteomics data generated using Trypsin/LysC digestion against a custom proteome database with Spectronaut (v20.3.251215). Protein intensities were imputed for protein groups using downshifted imputation in Perseus (82) (v1.5.15.0, width = 0.3, down shift = 1.8) if a protein was detected in at least two samples in at least one of the groups (FCX, CAUD, or CBM), and in fewer than two samples in the remaining groups. A combined quantification matrix containing imputed protein groups was quantile normalised using the normalizeBetweenArrays utility of limma (v3.56.2) (83) and subjected to principal component analysis (PCA) to visualise the clustering of samples within each group. Differential expression analysis was performed on the normalised intensity values using limma. A design matrix was constructed based on brain region (FCX, CAUD, CBM), and a linear model was fit to each protein group using lmFit. Pairwise contrasts (CBM vs. CAUD, CBM vs. FCX, and CAUD vs. FCX) were extracted using contrasts.fit, and moderated t-statistics were computed using eBayes. Protein groups with a log_2_ fold change ≥1.5 and Benjamini-Hochberg adjusted *P* value <0.05 were considered differentially regulated.

#### Integration of melanoma variant and peptide data

Genomic coordinates of variants were intersected with peptide genomic coordinates to assign each variant to a peptide. Variants detected in at least two of the three resistant samples were considered for further analysis. The median allele frequency was calculated for the resistant and pre- and early-treatment groups, and the difference (resistant minus pre andearly) was computed. Variants showing a positive difference were considered enriched in the resistant group. Genes from variant peptides with a CADD score >10 and showing a 10% increase in both allele frequency and variant peptide abundance in the resistance group were considered for pathway enrichment analysis.

Gene-level expression estimates derived from RNA-seq data were used to infer transcription factor (TF) activity with the decoupleR R package, applying the univariate linear model (ULM) method (84) and the DoRothEA regulon as the prior gene regulatory network (85).

## Data visualisation

Data processing and visualisation were performed using R packages, including rtracklayer (86), GenomicRanges (87), and the tidyverse suite.

## Acknowledgements

We thank Nicholas Williamson, Ching-Seng Ang, Shuai Nie, Swati Varshney and Michael Leeming for instrument support in the Bio21 Mass Spectrometry and Proteomics Facility. This project was supported by the Department of Anatomy and Physiology Flagship Research Program Initiative to C.A.W, M.B.C and B.L.P, and a School of Biomedical Sciences EMCRA award to H.K. from the University of Melbourne. We would like to acknowledge that brain tissues were received from the VBB, supported by The Florey, The Alfred, and the Victorian Institute of Forensic Medicine and funded in part by Parkinson’s Victoria, MND Victoria, and FightMND. We further acknowledge the donors and their families for the selfless donations to research. We gratefully acknowledge the Melbourne Research Cloud for providing infrastructure support for the GenomeProt webserver. We also thank Dr. Linda Nguyen and the ADAPT Lab team at the University of Melbourne for their assistance in designing the GenomeProt logo. The melanoma xenograft studies were supported by a Victorian Cancer Agency Mid-Career Fellowship awarded to L.S. and we gratefully acknowledge the Cancer Models Translational Research Centre for mouse husbandry and the Molecular Genomics Core facility for genomic DNA sequencing at the Peter MacCallum Cancer Centre. This work was supported by NHMRC Investigator Grants to M.B.C. (GNT1196841) and B.L.P (GNT2009642).

## AUTHOR CONTRIBUTIONS

M.B.C. and B.L.P. conceptualised the study. R.D.P.-I., M.D., A.A., L.S., and B.L.P. performed experiments and generated the data. H.K., J.G., C.Y.W., R.D.P.-I., A.L., L.S. and B.L.P. analysed the data. Y.D.J.P led software testing. H.K., J.G., and C.Y.W. performed bioinformatics and built GenomeProt. H.K., C.A.W., L.S., M.B.C., and B.L.P. provided resources, supervised and funded the research. H.K., M.B.C., C.A.W., and B.L.P wrote the manuscript. All authors read and approved the final manuscript.

## ETHICS APPROVAL AND CONSENT TO PARTICIPATE

Control post-mortem human brain samples were obtained from five consented individuals collected by the Victorian Brain Bank (VBB). Ethical approval was provided by the Human Research Ethics Committee of the University of Melbourne: #12457 and #28304.

## DECLARTION OF INTERESTS

J.G., R.D.P., Y.D.J.P, L.S., and M.B.C. have received support from Oxford Nanopore Technologies (ONT) to present their findings at scientific conferences. However, ONT played no role in the study design, execution, analysis, or publication of this research.

## Data and Code Availability

The GenomeProt code is available at https://github.com/ClarkLaboratory/GenomeProt.git. The GenomeProt web server is accessible at https://genomeprot.researchsoftware.unimelb.edu.au.

- The Kusa long-read RNA-seq and Melanoma WGS and RNA-seq raw sequencing data have been deposited in the Sequence Read Archive (SRA) under the BioProject ID PRJNA1474681.
- The proteogenomics data generated in this study are deposited to the ProteomeXchange Consortium (http://proteomecentral.proteomexchange.org/cgi/GetDataset) via the PRIDE (88) and can be accessed in the following projects:

- Proteogenomics analysis of Kusa 4B10 Cells: Project accession: PXD079484; Token: qxVOEEtUurVm
- Proteogenomics analysis of human brain: Project accession: PXD080085; Token: 0PR9wDSnVJuR
- Proteogenomic analysis of melanoma resistance: Project accession: PXD080265; Token: 1wVRHDcsVqDq
- Synthetic peptide data can be accessed via Panorama Web Repository (89) through the accession code Panorama Web: U of Melbourne – Parker Lab: Novel synthetic peptides in human brain
- Human postmortem brain whole transcriptome, long-read (ONT) data have been deposited in the European Genome-phenome Archive (EGA) and are available for download at EGA (EGAS00001008495). In accordance with donor consent and ethical approvals, the files can be provided pending scientific review and a completed data access agreement.

## Supplementary Table Legends

**Table S1-** GenomeProt features and usability compared to other recent proteogenomic pipelines with a specific focus on long-read RNA-seq.

**Table S2-** Proteogenomic analysis of Kusa 4B10 cells using GenomeProt.

**Table S3-** Splicing analysis of Kusa 4B10 using both long-read RNA-seq and proteomics data.

**Table S4-** Transcriptomic analysis of human brain regions using long-read RNA-seq.

**Table S5**- Peptides identified from ncORFs via proteogenomic analysis using GenomeProt in the human brain.

**Table S6**- *In silico* characteristics of ncORF identified in the human brain.

**Table S7**- Synthetic peptide analysis of ncORFs identified in the human brain.

**Table S8**- Transcriptomic analysis of melanoma using long-read RNA-seq.

**Table S9**- Variant peptides identified by a proteogenomic analysis using GenomeProt in melanoma.

**Table S10**- Variant peptides with increased allele frequency and abundance during treatment resistance.

## Notes

### Competing Interest Statement

J.G., R.D.P., L.S., and M.B.C. have received support from Oxford Nanopore Technologies (ONT) to present their findings at scientific conferences. However, ONT played no role in the study design, execution, analysis, or publication of this research.

