## Supplementary Figures for "GenomeProt: User friendly proteogenomics for canonical and non-canonical proteoform characterisation"

* co-lead

Correspondence:

**Contents**

**p2-3: Supplementary figure 1. Database comparison.**

**p4: Supplementary Figure 2. Quality control and proteomic comparison of brain regions.**

**p5: Supplementary Figure 3. Sequence and evolutionary features of noncanonical ORFs (ncORFs) identified with proteomic evidence.**

**p6-11: Supplementary Figure 4. Validation of ncORF identified in human brain via comparison to synthetic peptides.**

**p12: Supplementary Figure 5. Quality control and proteomic comparison of resistance vs Pre-/Early-Treatment.**

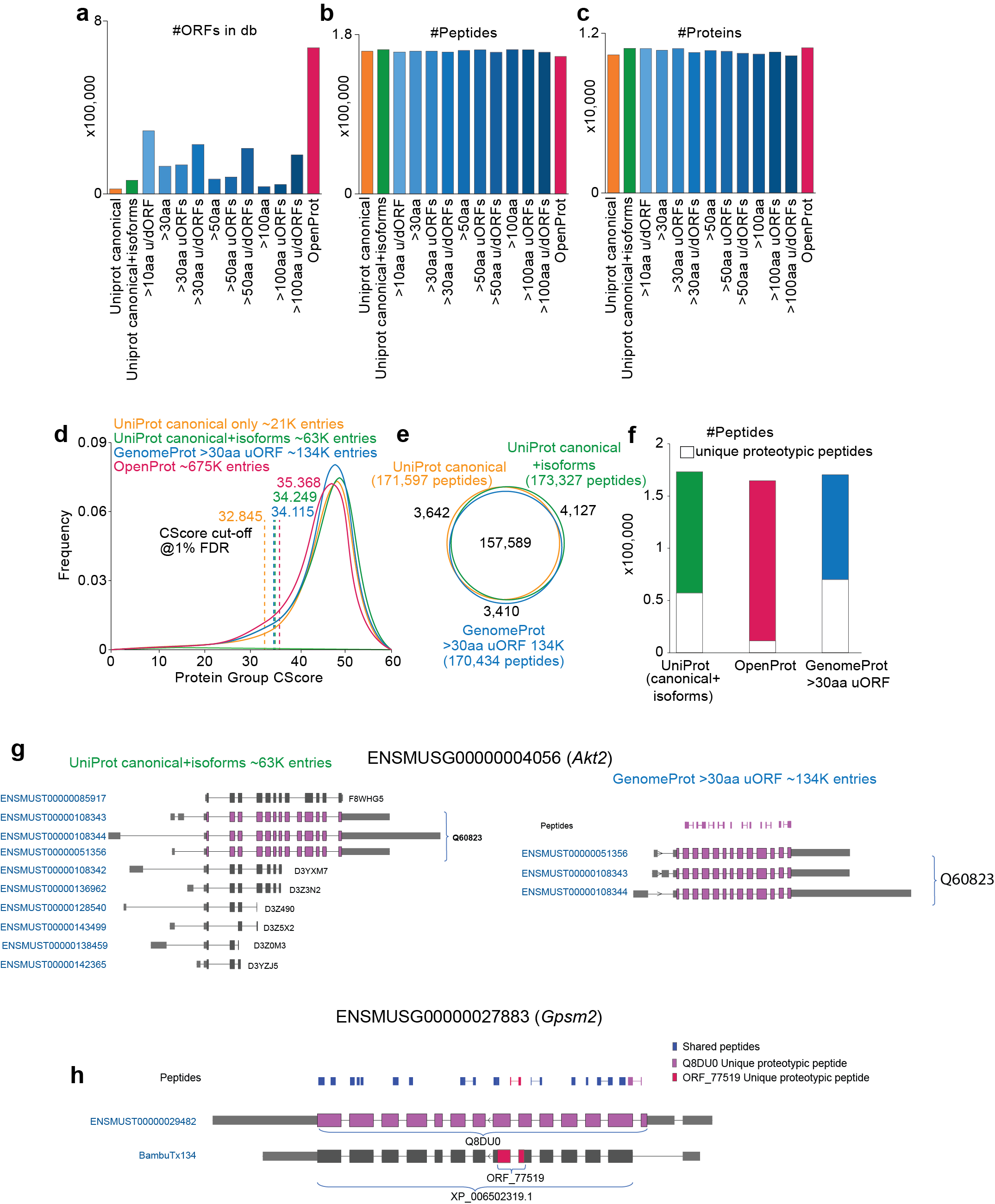

**Supplementary figure 1. Database comparison.** “>10aa u/dORF”: all ORFs greater than 10 amino acids (aa) and includes uORF and dORF; " >30aa”: all ORFs greater than 30aa but does not include uORF or dORF. “>30aa uORFs”: all ORFs greater than 30aa and includes uORFs greater than 10aa but does not include dORFs; “>30aa u/dORFs”: all ORFs greater than 30aa and includes both uORFs and dORFs greater than 10aa; " >50aa”: all ORFs greater than 50aa but does not include uORF or dORF; “>50aa uORFs”: all ORFs greater than 50aa and includes uORFs greater than 10aa but does not include dORFs; “>50aa u/dORFs”: all ORFs greater than 50aa and includes both uORFs and dORFs greater than 10aa; " >100aa”: all ORFs greater than 100aa but does not include uORF or dORF. “>100aa uORFs”: all ORFs greater than 100aa and includes uORFs greater than 10aa but does not include dORFs; “>100aa u/dORFs”: all ORFs greater than 100aa and includes both uORFs and dORFs greater than 10aa. **a**) Number of entries in each database tested. **b**) Number of peptides, and **c**) number of proteins identified in each database tested. **d**) Protein Group CScore distribution and cut-off at 1% false discovery rate (FDR) for each of the four databases listed. **e**) Venn diagram showing the overlap of identified peptides. **f**) Number of peptides identified and unique proteotypic peptides for each of the databases listed. **g**) Example of various proteoform entries of *Akt2* in the UniProt and GenomeProt databases and identified peptides. Proteoform found to be translated by GenomeProt database highlighted in purple. **h**) Peptides identified for *Gpsm2*. Highlighted in purple and red are the proteotypic peptides that uniquely map to the known and novel proteoforms encoded by a known and novel RNA respectively.

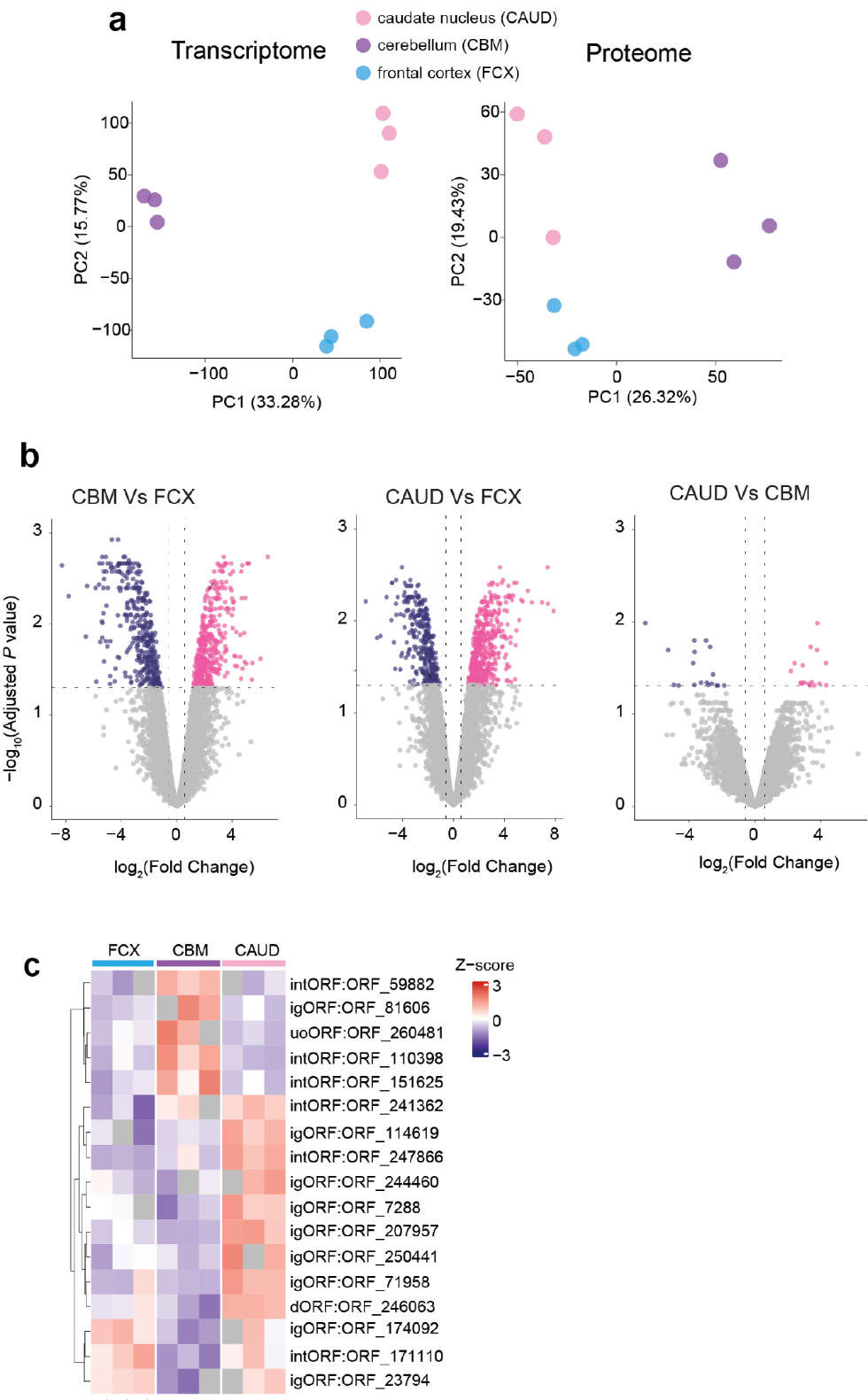

**Supplementary Figure 2. Quality control and proteomic comparison of brain regions.** (**a**) Principal component analysis of the transcriptome and proteome highlighting separation of the brain regions. (**b**) Volcano plot of the proteomic data showing differential abundance of proteins between the brain regions. (c) Differentially regulated ncORFs between brain regions.

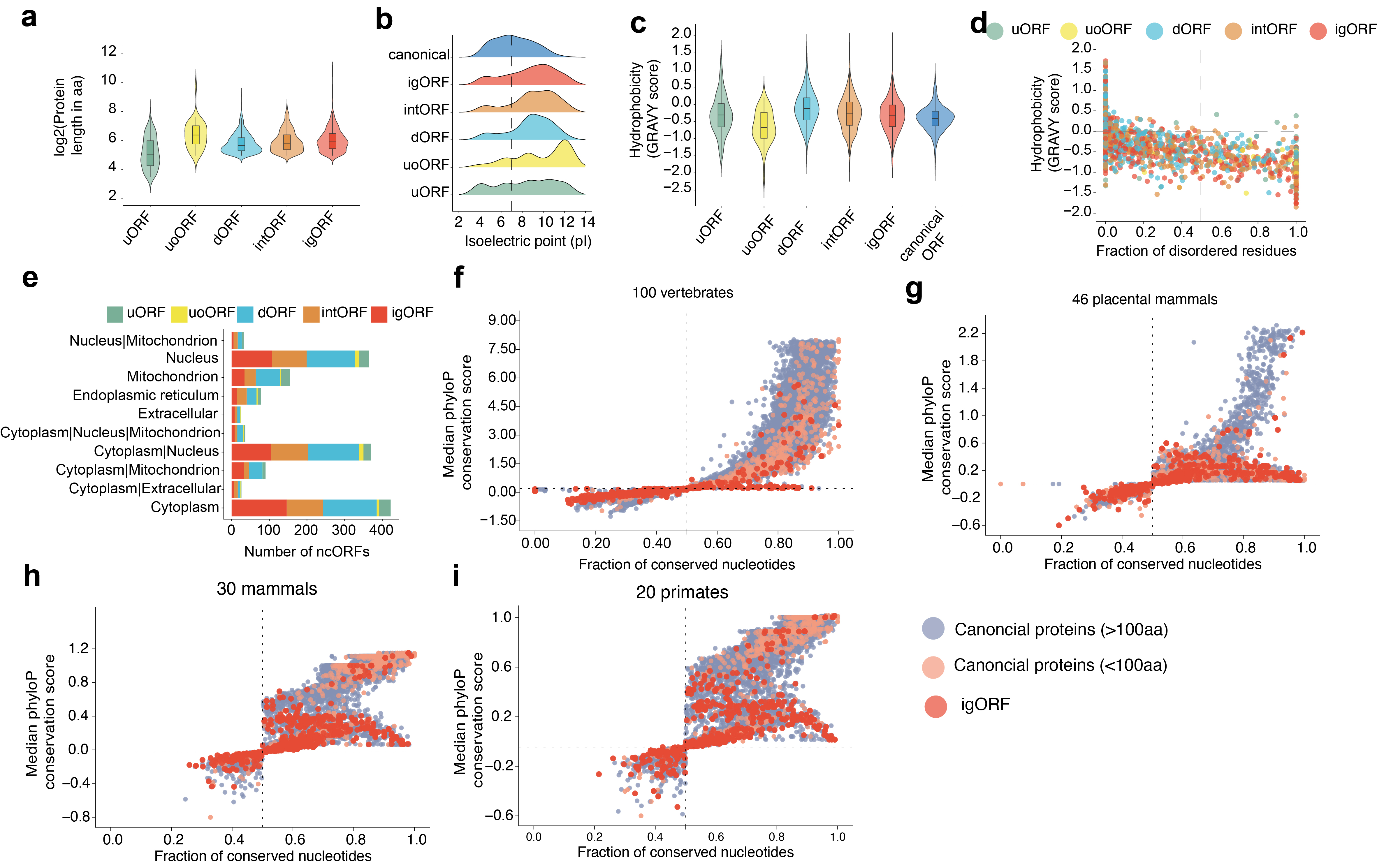
**Supplementary Figure 3. Sequence and evolutionary features of noncanonical ORFs (ncORFs) identified with proteomic evidence.** **(a)** Protein length distribution by ncORFs categories (uORF, uoORF, dORF, intORF, igORF). **(b)** Isoelectric point (pI) distribution by ncORF categories. The dotted vertical line indicates pH 7 (neutral pH). **(c)** Hydrophobicity (GRAVY score) distribution of ncORFs compared to canonical proteins. **(d)** Relationship between hydrophobicity (GRAVY score) and the fraction of disordered residues per ORF, colored by ORF categories (uORF, uoORF, dORF, intORF, igORF). Residues with a disorder score above 0.5 were considered disordered, and the fraction of disordered residues per ORF was calculated as the number of disordered residues divided by the total number of residues in the ORF. **(e)** Predicted subcellular localization of ncORFs, determined using DeepLoc 2.1, shown as the number of ncORFs assigned to each localisation category (or combination of categories) and colored by ORF categories. **(f–i)** Relationship between per-ORF median phyloP conservation score and the fraction of conserved nucleotides within the ORF across canonical proteins (>100 aa), canonical proteins (<100 aa), and igORFs, evaluated across four phylogenetic depths: 100 vertebrates (f), 46 placental mammals (g), 30 mammals (h), and 20 primates (i). The fraction of conserved nucleotides was calculated as the proportion of nucleotides within the CDS exceeding a phyloP score threshold of 0.2 (100 vertebrates) or 0 (46 placental mammals, 30 mammals, 20 primates) relative to the total number of nucleotides in the CDS.

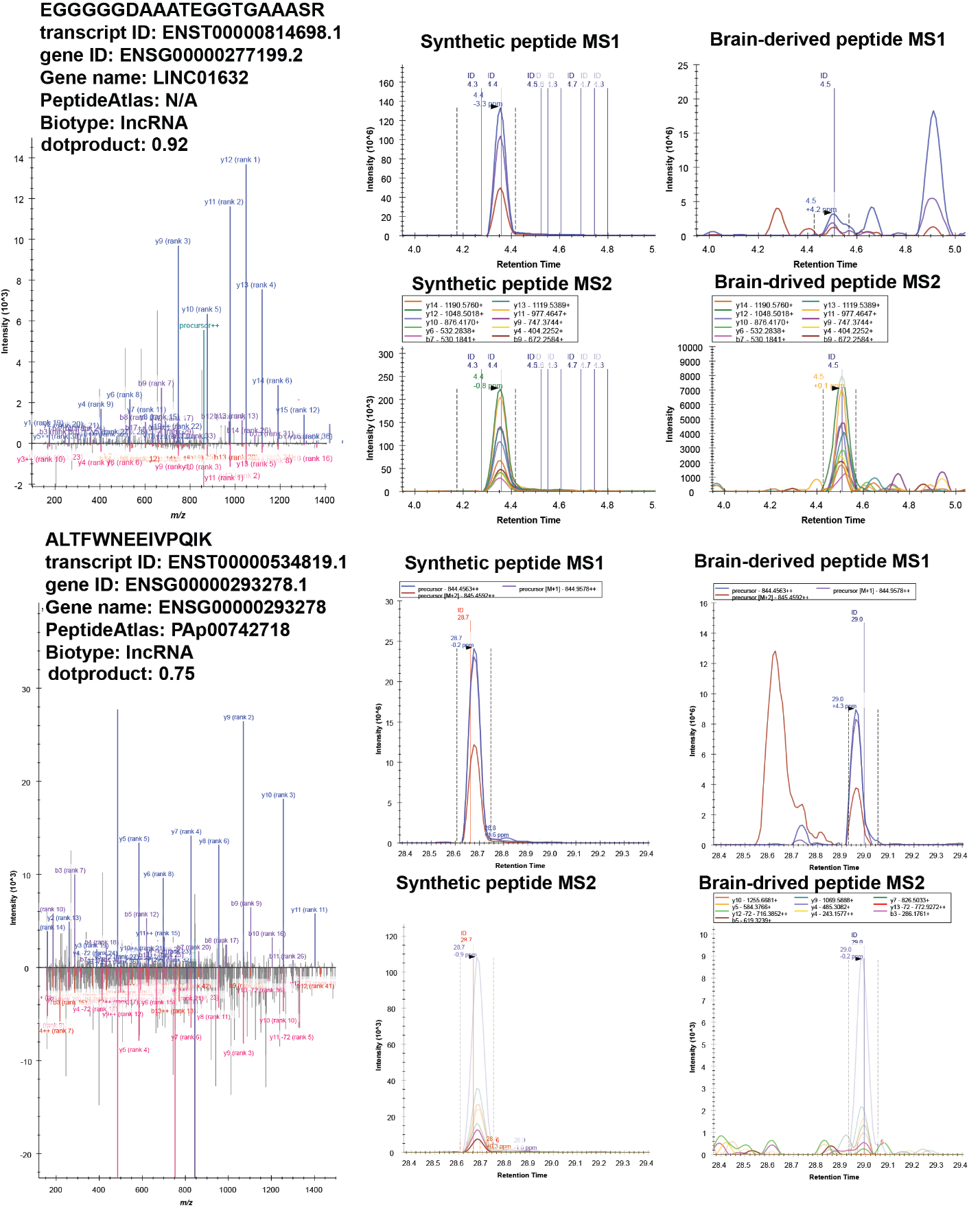

**b**

**a**

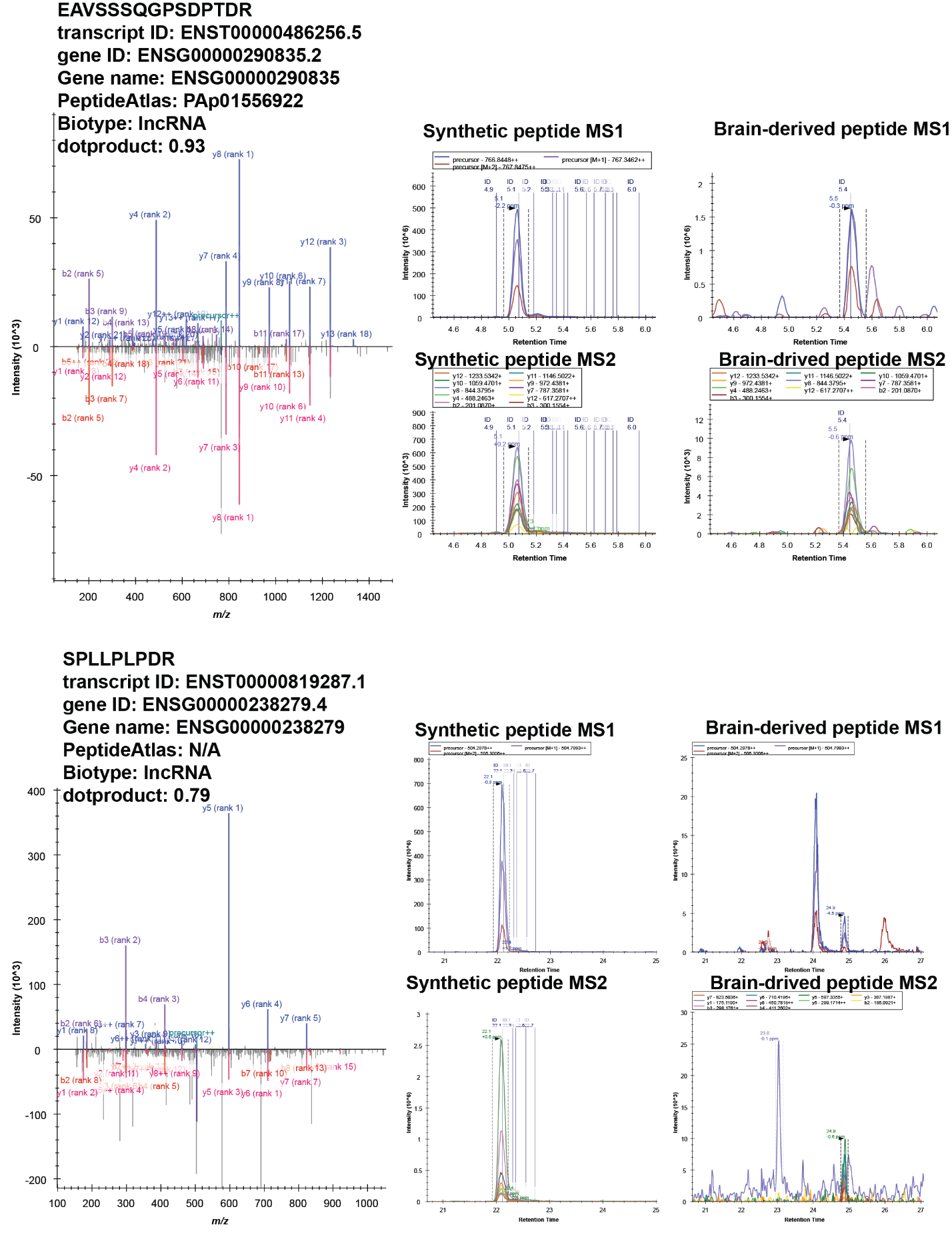

**d**

**c**

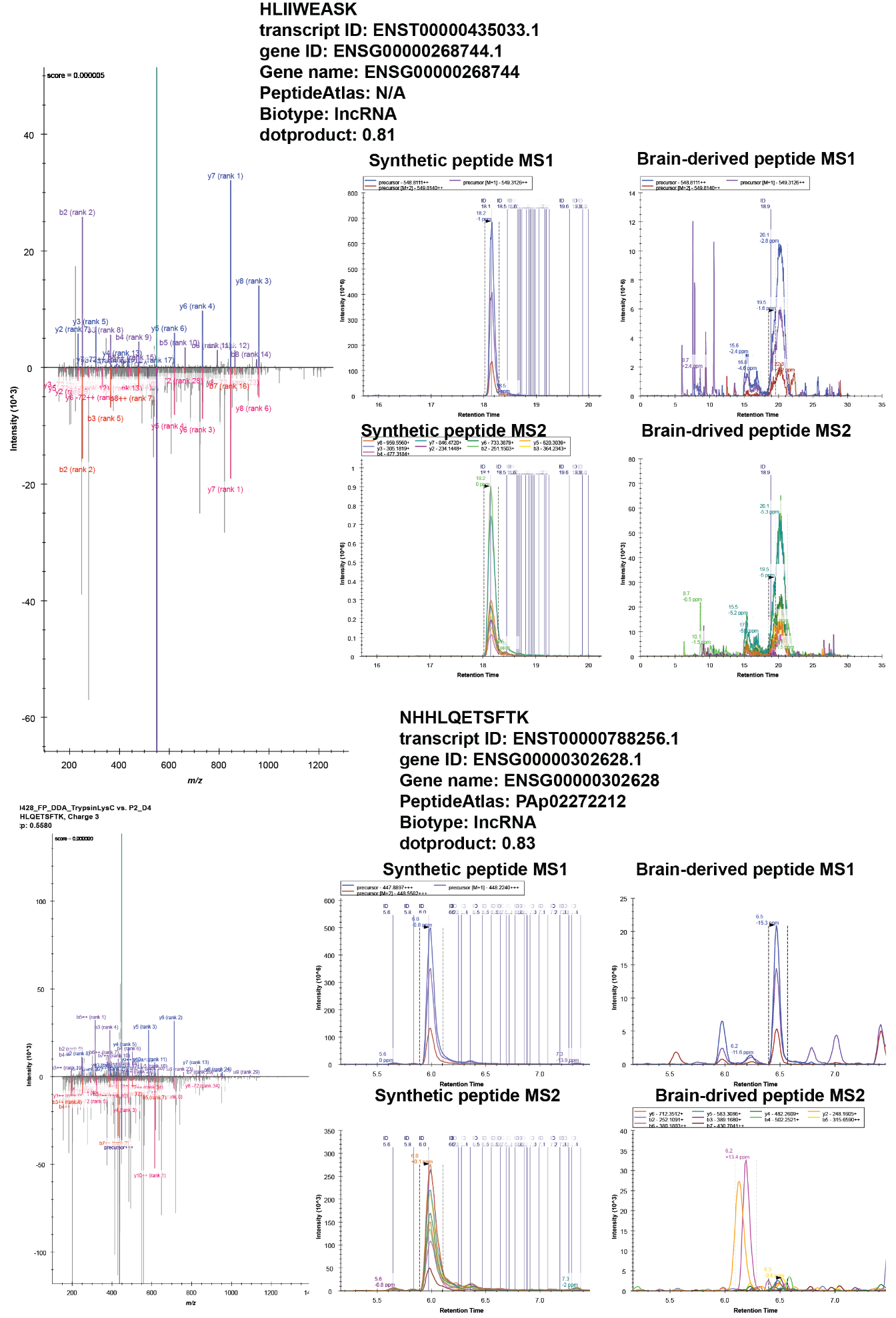

**f**

**e**

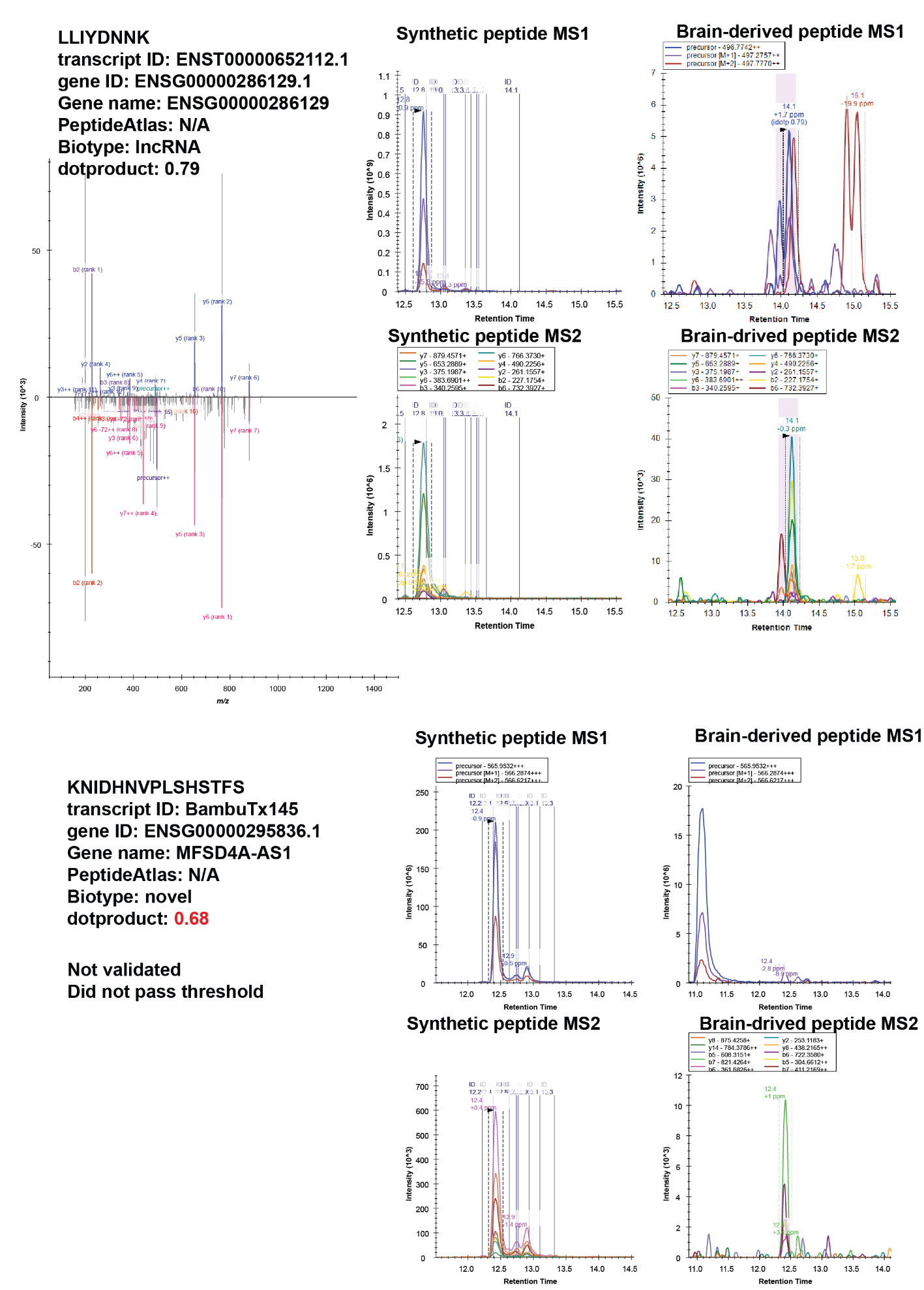

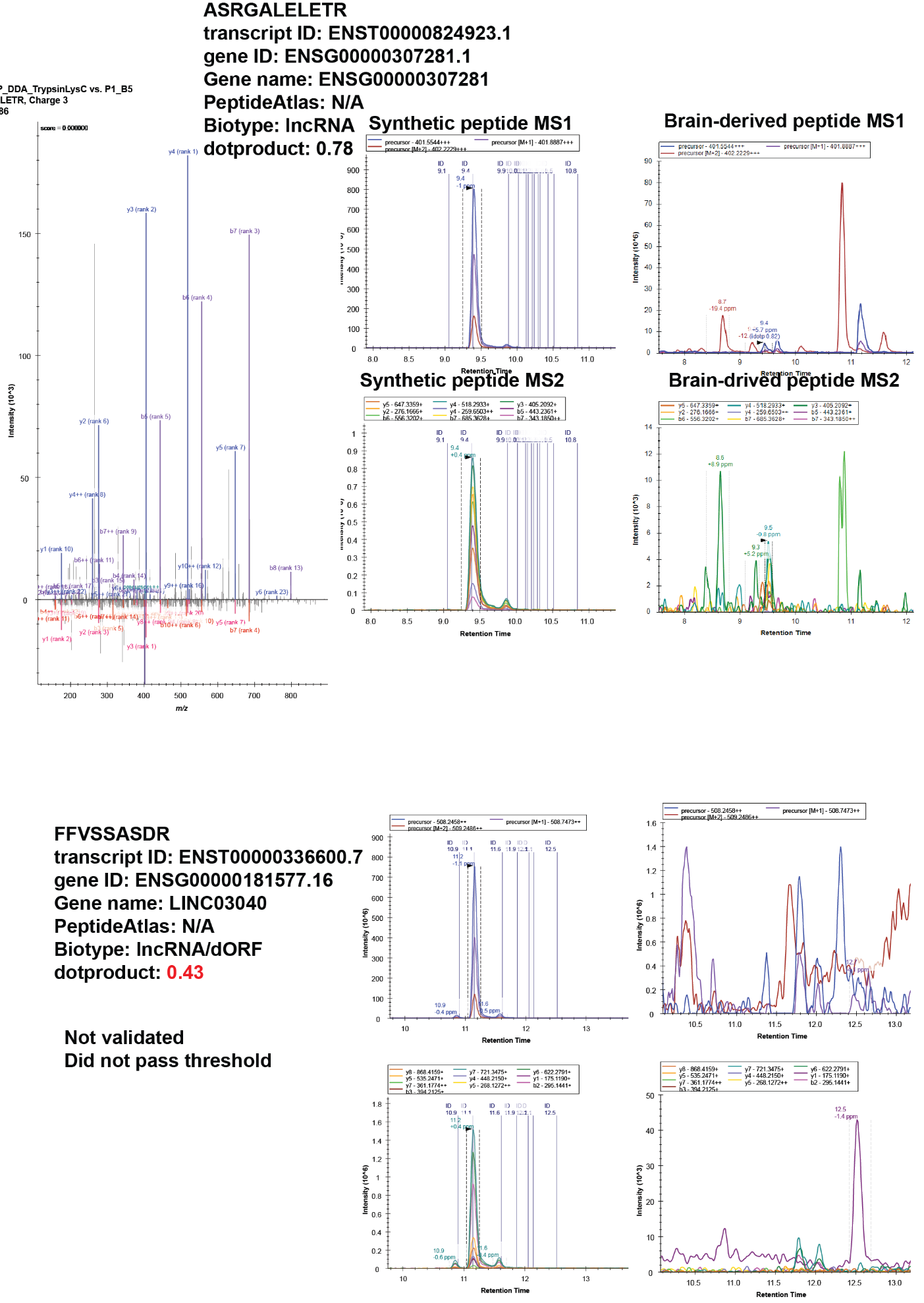

**g**

**h**

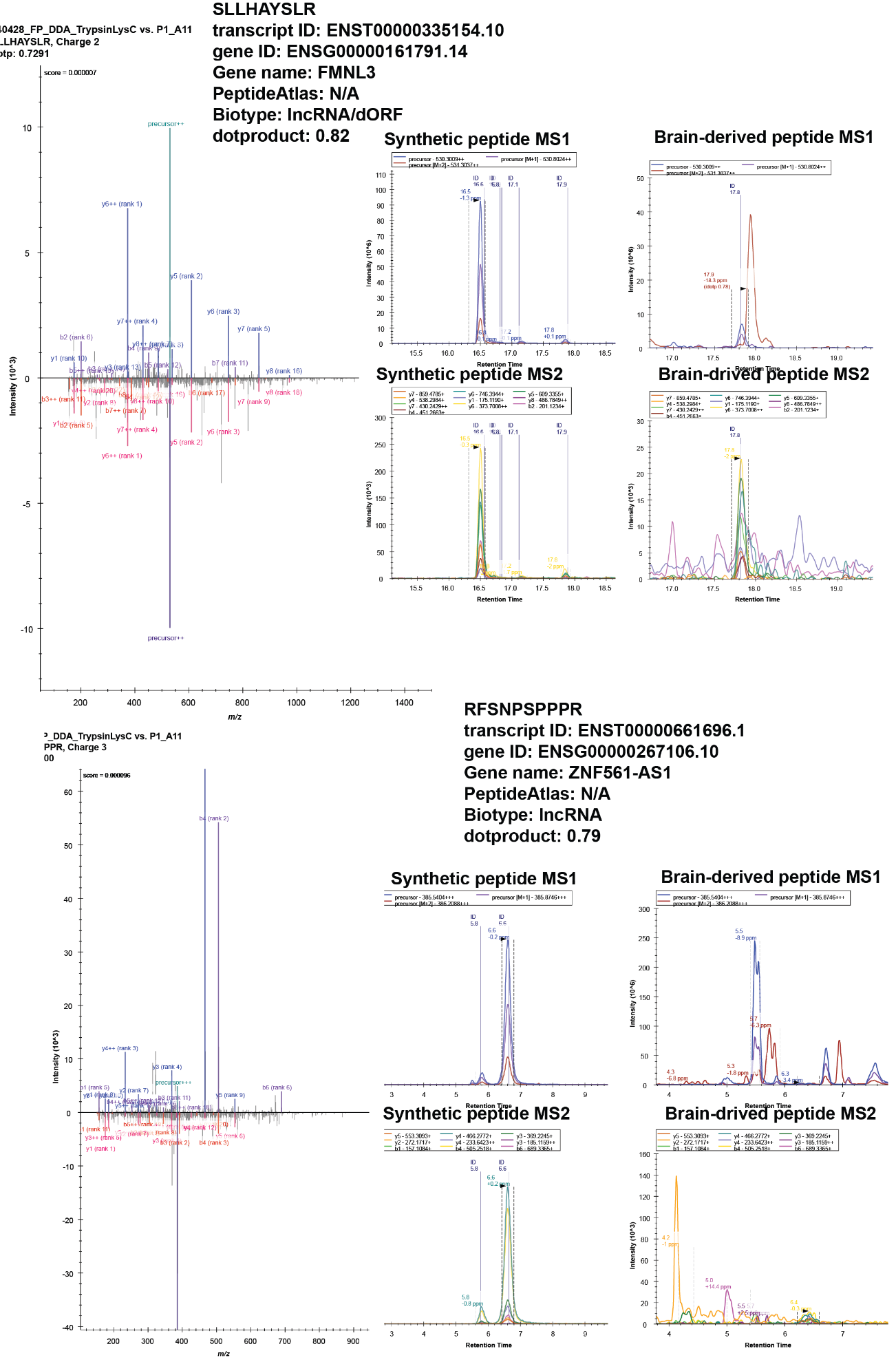

**i**

**j**

**Supplementary Figure 4. Validation of ncORF identified in human brain via comparison to synthetic peptides.** Left: Mirror MS2 spectra of synthetic peptide (top) and endogenous brain peptide (bottom). Right MS1 and MS2 extracted ion chromatograms. A total of ten peptides were validated: (**a**) EGGGGGDAAATEGGTGAAASR, (**b**) ALTFWNEEIVPQIK, (**c**) EAVSSSQGPSDPTDR, (**d**) SPLLPLPDR, (**e**) HLIIWEASK, (**f**) NHHLQETSFTK, (**g**) ASRGALELETR, (**h**) LLIYDNNK, (**i**) SLLHAYSLR, (**j**) RFSNPSPPPR.

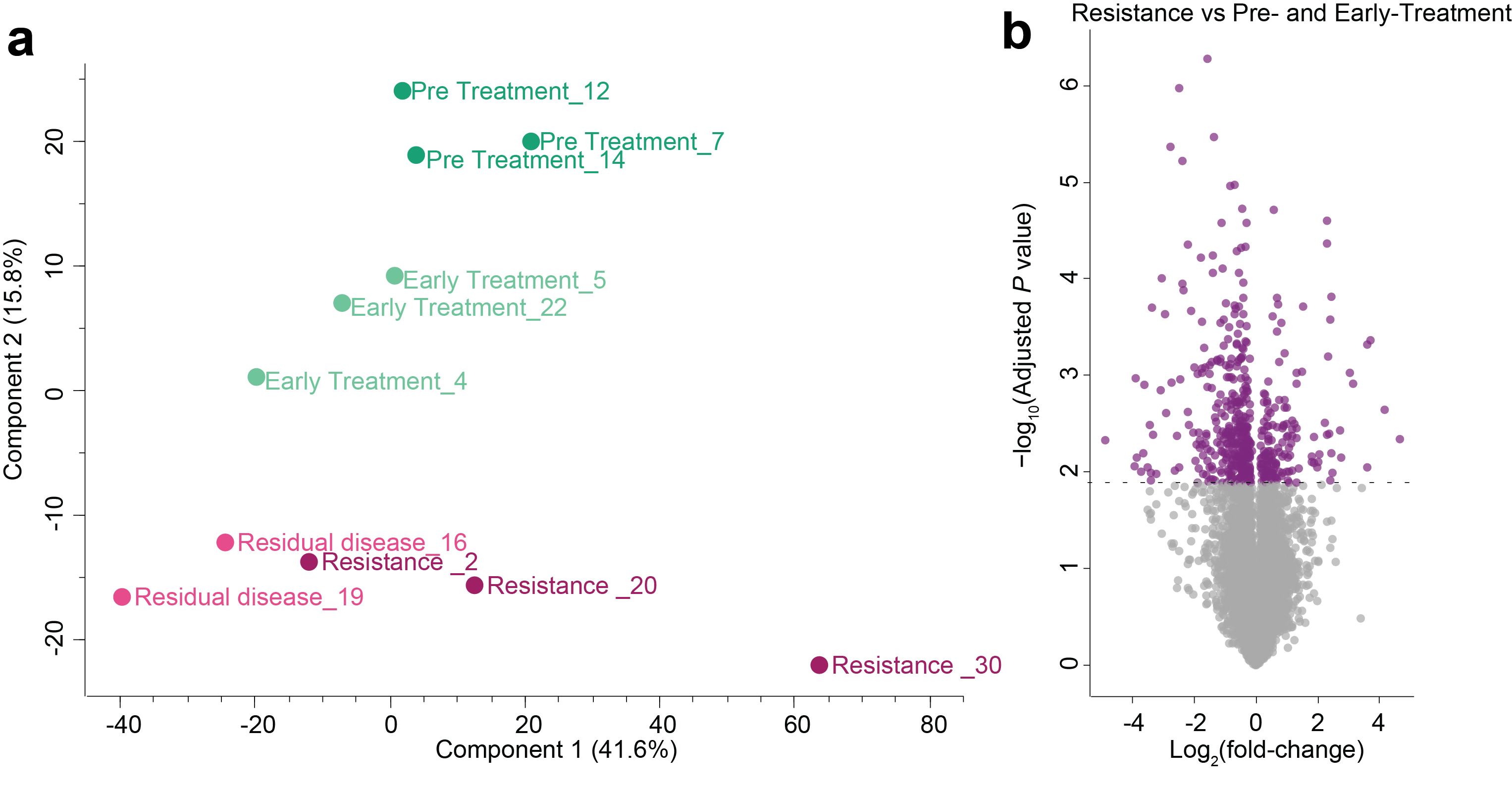

**Supplementary Figure 5. Quality control and proteomic comparison of resistance vs Pre- and Early-Treatment. a**) Principal component analysis of the proteomic data. **b**) Volcano plot of regulated proteins in the resistance group vs the Pre- andearly-treatment group.
