## Supplementary Tables for "GenomeProt: User friendly proteogenomics for canonical and non-canonical proteoform characterisation": Table_S1.docx

**Table S1- GenomeProt features and usability compared to other recent proteogenomic pipelines with a specific focus on long-read RNA-seq.**

| **Features** | **ProteomeGenerator3 (PG3)** | **Miller et al *Genome Biol* 2022** | **Schertzer et al *bioRxiv* 2026** | **GenomeProt** |
| --- | --- | --- | --- | --- |
| Reference | [DOI link](https://doi.org/10.7554/eLife.108878.1) | [DOI link](https://doi.org/10.1186/s13059-022-02624-y) | [DOI link](https://doi.org/10.64898/2026.05.27.728216) | N/A |
| Graphical user interface (GUI) | No | No | No | Yes (R Shiny application) |
| User modes | Command-line | Command-line | Command-line | GUI, command-line and public server |
| Installation and dependencies | Docker and Singularity | Docker and Conda environment | Apptainer, Singularity, Docker and Conda environment | Docker and Conda environment |
| Parallel execution | Yes (Nextflow) | Yes (Nextflow) | Yes (Nextflow) | No |
| Main input data | - Oxford Nanopore RNA-seq reads (BAM file) | - PacBio RNA-seq reads (BAM) - Mass spectrometry peptide data | - PacBio RNA-seq reads (BAM or FASTQ) - Mass spectrometry peptide data | - Oxford Nanopore, PacBio or Illumina RNA-seq reads (FASTQ, BAM, GTF) - Mass spectrometry peptide data |
| Multi-species support | Yes | Human | Human and Mouse | Human, Mouse, Rat, Zebrafish, C. elegans and Fruit fly |
| Main output data | - FASTA | - FASTA - A TSV file listing details of the best predicted ORF for each PacBio accession - A TSV file containing classifications of all PacBio proteins - Peptide and protein BED files for creating visualization tracks in the UCSC Genome Browser | - FASTA - A TXT table containing classifications of all PacBio proteins - A summary TSV file of the annotated and novel peptides that are mapped to transcripts - Peptide and protein BED files for creating visualization tracks in external visualizers, such as IGV and the UCSC Genome Browser | - FASTA - A TSV file with details of predicted and annotated proteins in the database - A proteogenomics integration summary HTML report and a TSV file with mapped peptides and transcript annotations - A visualizable, comprehensive GTF annotation file on the peptides, ORFs and transcripts identified, all mapped to genomic coordinates - Peptide, ORF and transcript BED files for creating visualization tracks in the UCSC Genome Browser |
| Steps carried out | - Proteome database generation | - Proteome database generation - Peptide search - Proteogenomics data integration | - Proteome database generation - Peptide search - Proteogenomics data integration | - Proteome database generation - Peptide search - Proteogenomics data integration - Data visualization |
| Sequencing data input compatibilities | - Optimized for long-read data (BAM) | - Long-read PacBio CCS (circular consensus sequencing) data (BAM) | - Post-processed (i.e. have already undergone deconcatenation, demultiplexing and primer removal) PacBio FLNC (full-length non-chimeric) read data (FASTQ and BAM) | - Distinct workflows for short-read and long-read data (FASTQ, BAM or GTF) |
| ORF database annotation | - UniProt-style headers - Includes unannotated proteins with nomenclature: (“PG” + protein index) - Limited ORF annotations: ORF accession, transcript ID, Ensembl / Bambu gene ID, UniProt match status | - UniProt-style headers - Includes unannotated proteins formatted as PacBio accessions (“PB.” + gene index number + “.” + isoform index number) - Comprehensive ORF annotations but spread across many output files: ORF accession, transcript ID, gene ID and symbol, strand, SQANTI transcript and protein biotypes, ORF genomic coordinates, reading frame, ORF classification (full splice match (FSM) / incomplete splice match (ISM) / novel in catalog (NIC) / novel not in catalog (NNC), each with subcategories), CPAT ORF coding probability, Fickett TestCode score, Hexamer score, ATG start codon score, ORF calling confidence, CPM values, UniProt match status, protein sequence | - UniProt-style headers - Includes unannotated proteins formatted as PacBio accessions (“PB.” + gene index number + “.” + isoform index number) and/or deterministic SHAKE256 hashes based on the splice junction chain of the transcript - Comprehensive ORF annotations but spread across many output files: ORF accession, transcript ID, gene ID and symbol, strand, SQANTI transcript and protein biotypes, ORF genomic coordinates, reading frame, ORF classification (FPM (full protein match) / IPM (incomplete protein match) / NPC (novel protein combination) / NPE (novel protein element) / intergenic / genic / antisense / fusion / orphan FSM (full splice match) / orphan ISM (incomplete splice match) / orphan NIC (novel in catalog) / orphan NNC (novel not in catalog) / orphan monoexon, each with subcategories), CPAT ORF coding probability, ORF calling confidence, CPM values, PSM (peptide spectrum match) scores, UniProt match status, protein sequence, peptide intensities | - UniProt-style headers - Includes unannotated proteins with nomenclature: (“ORF_” + Protein index) - Comprehensive ORF annotations in a single file, including: ORF accession, gene ID and symbol, protein functional annotation (if available), transcript ID, strand, transcript biotype, transcript coordinates, ORF genomic coordinates, reading frame, ORF classification based on genomic/transcript location, OpenProt ID, protein sequence, amino acid change (for variant proteins), longest ORF status within the transcript, UniProt status (reviewed/unreviewed), and physicochemical properties such as molecular weight, isoelectric point, hydrophobicity, and aliphatic index |
| Variant-aware proteome database generation | Variants included only if present in assembled transcript sequences; no direct VCF input | Variants included only if present in assembled transcript sequences; no direct VCF input | Variants included only if present in assembled transcript sequences; no direct VCF input | - Incorporates Single nucleotide variants (SNVs) from a multi-sample VCF file into the genome and generates variant-aware proteome databases. - Amino acid changes included in annotation files. |
| Peptide-to-genome mapping | Not supported | - Generates GTF and BED12 files showing the mapping of peptides onto genomic coordinates for visualization in the UCSC Genome Browser. - Includes peptide sequences; the most frequent amino acid preceding each peptide sequence; and the most frequent amino acid following each peptide sequence (taken from MetaMorpheus). | - Generates GTF and BED12 files showing the mapping of peptides onto genomic coordinates for use in external visualizers, such as IGV and the UCSC Genome Browser. - Includes peptide sequences and PSM scores. | - Generates a BED file showing the mapping of peptides onto genomic coordinates, which includes peptide sequences and the transcript IDs each peptide maps to. - Generates a combined annotations GTF file containing the information from the aforementioned BED file, in addition to the IDs and symbols of the genes each ORF is encoded in; ORF genomic coordinates; accessions of the ORFs a peptide is mapped to; type of peptide mapping (exonic or exon-spanning); whether a peptide uniquely maps to an ORF, gene or transcript; ORF localisation; transcript biotype; whether an ORF is the longest one in a transcript; and the UniProt status of the ORF (reviewed/unreviewed). - Generates a detailed peptide annotations TSV file containing most of the relevant information from the aforementioned GTF file, in addition to information such as overlapping exon coordinates, exon numbers, and physicochemical properties of ORFs (molecular weight, isoelectric point, hydrophobicity and aliphatic index). |
| Proteins / ORFs identified and genomic annotation | - Not supported | - Generates GTF and BED12 files showing the mapping of proteins onto genomic coordinates for visualization in the UCSC Genome Browser. - Includes protein classifications (FSM / ISM / NIC / NNC) and CPM values. | - Generates GTF and BED12 files showing the mapping of proteins onto genomic coordinates for use in external visualizers, such as IGV and the UCSC Genome Browser. - Includes protein classifications (FPM / IPM / NPC / NPE / intergenic / genic / antisense / fusion / orphan FSM / orphan ISM / orphan NIC / orphan NNC / orphan monoexon) and, for each ORF, ratios of the average ORF CPM to the total average ORF CPM of the gene it is encoded in. | - Generates a BED file showing the mapping of ORFs onto genomic coordinates, which includes ORF accessions and the transcript IDs each ORF is encoded in. - The aforementioned peptide annotations TSV file and combined annotations GTF file contain information on the ORFs that have peptide mappings. - Includes protein classifications (5’UTR [uORF], 5’UTR-CDS [uoORF], 3’UTR [dORF], intergenic [igORF], gene overlap [intORF]) |
| Peptide visualization | Not supported | Not supported (need to upload the relevant output files into an external visualizer, e.g. the UCSC Genome Browser) | Not supported (need to upload the relevant output files into an external visualizer, e.g. the UCSC Genome Browser) | - Interactive and customizable genome- and transcript-level visualization - Comparison of transcript and peptide abundances across samples - Downloadable plots in PDF, SVG, PNG and JPEG formats |
| Results | Generated in multiple directories | Generated in multiple directories | Generated in multiple directories | - If the GenomeProt website GUI is used, results are downloadable as compressed zip files - If GenomeProt is run on the command line, results are generated in user-specified directories |
